# Primary complex motor stereotypies are associated with de novo damaging DNA coding mutations that identify candidate risk genes and biological pathways

**DOI:** 10.1101/730952

**Authors:** Thomas V. Fernandez, Zsanett P. Williams, Tina Kline, Shreenath Rajendran, Farhan Augustine, Nicole Wright, Catherine A. W. Sullivan, Emily Olfson, Sarah B. Abdallah, Wenzhong Liu, Ellen J. Hoffman, Abha R. Gupta, Harvey S. Singer

**Author notes:** Corresponding author. Please address correspondence to: Thomas V. Fernandez, MD, Yale University School of Medicine 230 S Frontage Road, New Haven, CT 06520.

## Abstract

Motor stereotypies are common in children with autism spectrum disorder (ASD), intellectual disability, or sensory deprivation, as well as in typically developing children (“primary” stereotypies, CMS). The precise pathophysiological mechanism for motor stereotypies is unknown, although genetic etiologies have been suggested. In this study, we perform whole-exome DNA sequencing in 129 parent-child trios with CMS and 853 control trios (118 cases and 750 controls after quality control). We report an increased rate of de novo predicted-damaging variants in CMS versus controls, identifying *KDM5B* as a high-confidence risk gene and estimating 184 genes conferring risk. Genes harboring de novo damaging variants in CMS probands show significant overlap with those in Tourette syndrome, ASD candidate genes, and those in ASD probands with high stereotypy scores. Furthermore, exploratory biological pathway and gene ontology analysis highlight histone demethylation, organism development, cell motility, glucocorticoid receptor pathway, and ion channel transport. Continued sequencing of CMS trios will identify more risk genes and allow greater insights into biological mechanisms of stereotypies across diagnostic boundaries.

## INTRODUCTION

Motor stereotypies are rhythmic, repetitive, prolonged, fixed, patterned, non-goal directed movements that are often bilateral and temporarily stop with distraction. Complex motor stereotypies (CMS) include hand flapping, finger wiggling, head nodding, and rocking; these are often accompanied by mouth opening, head posturing, jumping, pacing and occasional vocalizations (1). Movements occur for up to minutes in duration, multiple times per day, and tend to be exacerbated by excitement, fatigue, stress, boredom, or being engrossed in an activity. CMS are common in children with autism spectrum disorder (ASD), intellectual disability, or sensory deprivation, as well as in typically developing children. A favored classification subdivides by etiology into primary (otherwise typically developing) and secondary categories. In both groups, stereotypies often result in social stigmatization, classroom disruption, and interference with academic activities.

In children with ASD, stereotypic behaviors (“secondary” stereotypies) occur in about 44% of patients and are recognized as a core phenotype of the disorder (2). The severity and frequency of motor stereotypies is correlated with severity of illness, degree of intellectual disability, and impairments in adaptive functioning and symbolic play (3–9). They are often associated with self-injurious behaviors (10, 11). A wide range of medications have been tried for treatment of stereotypies in ASD, but efficacy is inconsistent and inadequate, with potential for long-term side effects (12).

Motor stereotypies also occur in otherwise typically developing children (“primary” stereotypies) (13–22). Studies comparing primary and secondary stereotypies show that there is considerable similarity in their phenomenology (23–25). Primary CMS has a typical age of onset before 3 years, and greater than 90% of children continue to experience CMS into adolescence and adulthood (16, 26). The prevalence of primary CMS is estimated to be 3-4% of children in the U.S. (26, 27). Similar to secondary stereotypies, medications are generally regarded as ineffective for primary CMS (13, 28), but there is evidence to support the benefits of cognitive behavioral therapy (29, 30).

The precise pathophysiological mechanism for motor stereotypies remains obscure (31), though investigators have hypothesized abnormalities within cortico-striatal-thalamo-cortical pathways (32–38) and several neurotransmitter systems (33, 39–41). A genetic etiology for stereotypies has been suggested in primary and secondary categories, although the specific gene(s) contributing to this movement disorder remain unclear. With respect to secondary stereotypies in ASD, family studies have demonstrated that these repetitive behaviors are highly heritable, with a genetic etiology that is likely independent from other core diagnostic features (42). While there are no studies of recurrence risk or twin concordance reported for primary CMS, a positive family history is reported in 25-40%, while remaining cases appear to be sporadic (16, 28, 43).

Considering these findings and with the recent success of risk gene discovery in other neurodevelopmental disorders via detection of de novo, or spontaneous, germline DNA mutations (44–46), we conducted the first pilot genetic study of CMS in 129 typically-developing children and their parents. We hypothesized that primary CMS may represent a more genetically homogenous group of individuals versus those with secondary stereotypies, thereby facilitating genetic discovery and insight into the biology of stereotypies more generally (49, 50). Using whole-exome DNA sequencing, we identified an enrichment of de novo predicted-damaging mutations and identified one high-confidence risk gene, *Lysine Demethylase 5B* (*KDM5B*) in our cohort. By further analysis of de novo damaging mutations in primary CMS, we predict that there are approximately 184 CMS risk genes and that sequencing more CMS parent-child trios is a definite path toward discovering these genes. In this pilot study, we see an overlap between genes harboring de novo damaging mutations in CMS and those in Tourette syndrome. Furthermore, owing to the two de novo damaging *KDM5B* mutations in our CMS cohort, there is significant overlap with ASD probands with highest stereotypy scores, but not those with low scores. Finally, exploratory systems analyses of genes harboring de novo damaging mutations in CMS show enrichment in histone demethylation, multicellular organism development, and cell motility.

## MATERIALS AND METHODS

Figure 1 provides an overview of the study methods.

**Figure 1.**
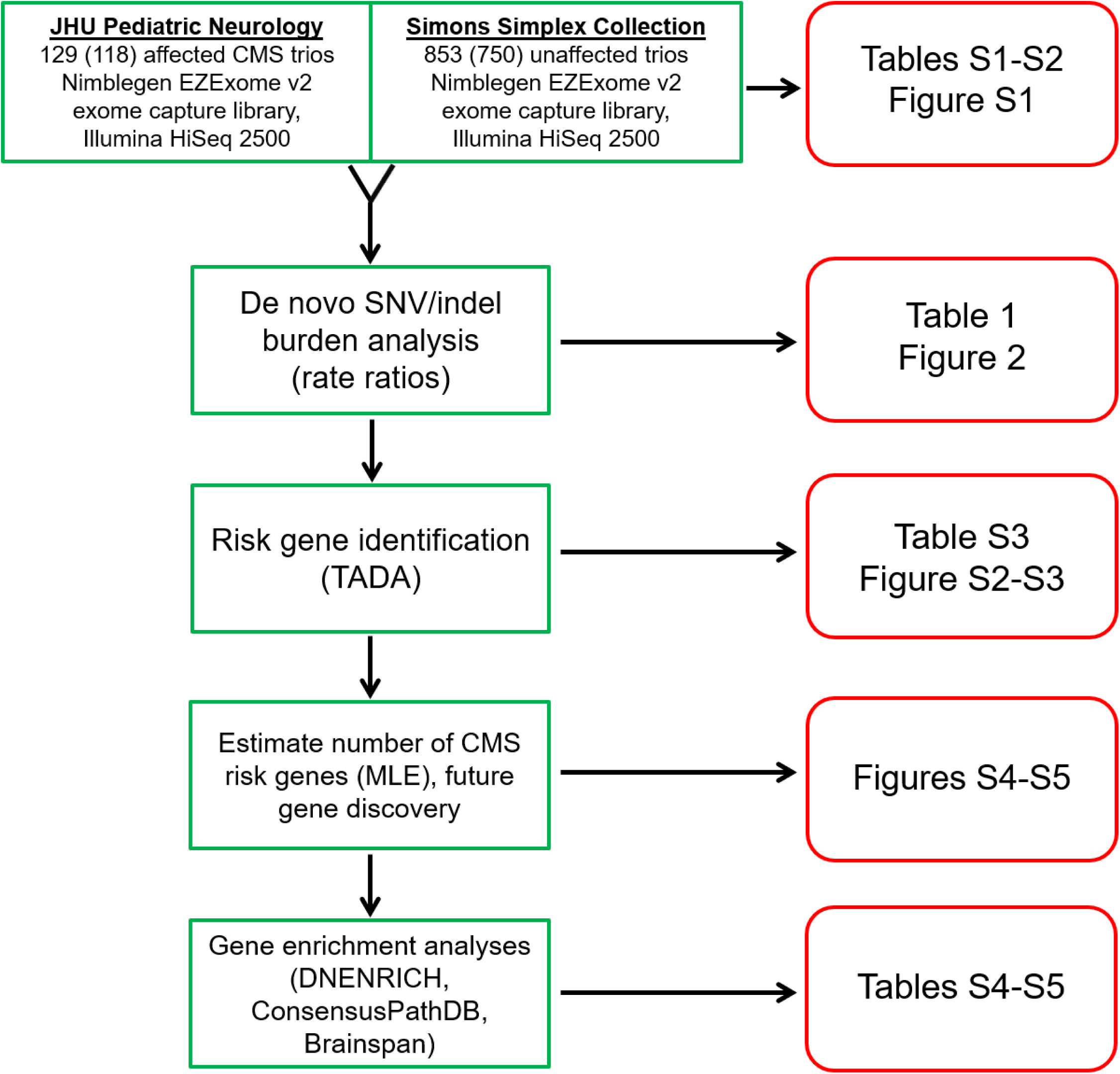
– Overview of variant discovery and data analysis. We performed whole exome sequencing on 129 CMS and 853 control parent-child trios. After quality control, 118 CMS and 750 control trios remained for subsequent analyses. We performed a burden analysis, comparing the rates of de novo single nucleotide (SNVs) and insertion-deletion (indel) variants between cases and controls. Next, we assessed the significance of gene-level recurrence of de novo damaging variants in our CMS group, identifying one high-confidence risk gene. Using the MLE method, variant simulations, and TADA, we estimated the number of genes contributing to CMS risk and used this estimate to predict the number of risk genes that will be discovered as more CMS trios are sequenced. Finally, exploratory gene enrichment analyses were performed, assessing degree of overlap with gene sets from other disorders, canonical pathways, gene ontologies, and expression pattern clustering within certain brain regions across development.

### Subjects and Assessment Measures

This protocol was approved by the Johns Hopkins Medicine Institutional Review Board. Children with primary complex motor stereotypies (CMS) were recruited from either the Johns Hopkins Pediatric Neurology Movement Disorder Outpatient Clinic (HSS, Director), or via email. All participants verbally consented and signed parental consent was obtained. Using standardized forms via telephone, the study coordinator completed a brief screening general history, obtained baseline data about each child’s stereotypies, and completed an Autism Spectrum Screening Questionnaire (ASSQ). The presence of stereotypic movements was confirmed, either via direct observation in clinic or by video review (HSS). If the subject passed the screening assessment, additional data was collected on the child and both parents via RedCap, an electronic web-based application for data capture and online questionnaires. The latter included the Stereotypy Severity Scale (Motor and Impairment scores) and comorbidity measures (Multidimensional Anxiety Scale for Children—MASC; ADHD-Rating Scale IV; Conner’s Parent Rating Scale—CPRS; Repetitive Behavior Scale-Revised—RBS-R; Children’s Yale-Brown Obsessive-Compulsive Scale—CYBOCS; and Social Responsiveness Scale—SRS) (see Supplementary Methods).

For this pilot study, we prioritized the study of “simplex” CMS (children without known family history of affected first or second-degree relatives) to increase the likelihood of detecting de novo sequence and structural variants. Eligibility required participants to have: (a) confirmed complex motor stereotypies; (b) onset before age 3 years; (c) temporary suspension of movements by an external stimulus or distraction. Exclusion criteria included: (a) a total score >13 on the ASSQ or a prior autism spectrum disorder diagnosis; (b) evidence of intellectual disability; (c) seizures or a known neurological disorder; and (d) the presence of motor/vocal tics. The presence of inattentiveness, hyperactivity, or impulsivity (i.e., ADHD symptoms) and/or obsessive-compulsive behaviors were not exclusionary.

### DNA whole-exome sequencing (WES)

DNA was collected from all children meeting eligibility criteria and from their parents, using the Oragene OG-500 collection kit and standard extraction protocols (DNA Genotek, Ottowa, Ontario, Canada). Exome capture and sequencing were performed at the Yale Center for Genome Analysis (YCGA), using the NimbleGen SeqCap EZExomeV2 capture library (Roche NimbleGen, Madison, WI, USA) and the Illumina HiSeq 2500 platform (Illumina, San Diego, CA, USA). WES data from 853 unaffected parent-child trios (2,559 samples total) were obtained from the Simons Simplex Collection via the NIH Data Archive (https://ndar.nih.gov/edit_collection.html?id=2042). These children and their parents have no evidence of autism spectrum or other neurodevelopmental disorders (47). The same exome capture and sequencing platforms were used for these control samples.

### Sequence alignment, variant calling, and quality control

Alignment and variant calling of the sequencing reads followed the latest Genome Analysis Toolkit (GATK) (48) Best Practices guidelines, as described previously (49). Variants were annotated using RefSeq hg19 gene definitions using ANNOVAR (50). Trios were omitted from downstream analyses if (a) genetic markers were not consistent with expected family relationships; (b) an excessive number of de novo variants were observed, or (c) if they were outliers in principal components analysis (see Supplementary Methods). De novo variants were called as previously described (49) and as detailed in Supplementary Methods.

### Mutation rate and gene recurrence

Within each cohort, we calculated the rate of de novo mutations per base pair, using methods previously described (49). We included only those de novo variants present with a frequency of <0.001 (0.1%) in the ExAC v0.3.1 database (51) and compared de novo mutation rates in cases versus controls using a one-tailed rate ratio test (Supplementary Methods).

As described in our previous WES studies (45, 46, 52), we used the Transmitted And De novo Association (TADA-Denovo) test as a statistical method for risk gene discovery based on gene-level recurrence of de novo mutations within the classes of variants that we found enriched in CMS (53, 54). This test generates random mutational data based on each gene’s specified mutation rate to determine null distributions, then calculates a p-value and a false discovery rate (FDR) q-value for each gene using a Bayesian “direct posterior approach.” A low q-value represents strong evidence for CMS association. See Supplementary Methods for details.

### Estimating the number of CMS risk genes

As described previously (45, 52), we used a maximum likelihood estimation (MLE) method (55) to estimate the number of genes contributing risk to CMS, based on the observed number of de novo damaging variants in our dataset. See Supplementary Methods for details of these calculations.

Next, we used previously described methods (45, 52) to predict the likely number of risk genes that will be discovered as additional CMS parent-child trios are sequenced by WES.

These predictions utilize the estimated number of CMS risk genes along with CMS de novo mutation rates observed in our study to perform mutation simulations, followed by TADA-Denovo testing (see Supplementary Methods).

### Gene set overlap

We used DNENRICH (56) (https://psychgen.u.hpc.mssm.edu/dnenrich/) to test whether genes harboring de novo damaging mutations in our CMS subjects were significantly enriched among several gene lists from the literature related to neuropsychiatric disorders, including autism (ASD), schizophrenia (SCZ), Tourette’s disorder (TD), obsessive-compulsive disorder (OCD), attention-deficit/hyperactivity disorder (ADHD), and intellectual disability (ID). Additionally, we were interested in the question of whether our CMS cohort share genes harboring de novo damaging mutations with ASD probands having high stereotypy scores. To approach this question, we assembled lists of genes harboring damaging de novo mutations in ASD probands from the Simons Simplex Collection (SSC) for whom stereotyped behavior scores (Stereotyped Behavior Score from the RBS-R, Repetitive Behavior Scale-Revised) were available. We looked for overlap between our CMS cohort and those SSC ASD probands with stereotypy scores in the 90^th^ percentile (high stereotypies) and those scoring in the 10^th^ percentile (low stereotypies). These gene lists are compiled in Table S4. Further details about gene list curation and DNENRICH methods can be found in Supplementary Methods.

### Exploratory pathway, gene ontology, and spatiotemporal analyses

To determine whether genes harboring de novo damaging variants in CMS may perform similar biological functions, we used the list of CMS genes harboring de novo damaging mutations to identify enriched pathways and gene ontologies using ConsensusPathDB (Release 34, 15.01.2019). This tool integrates human protein and genetic interaction networks from 32 databases and interactions curated from the literature (57).

Finally, using this same list of genes, we searched for possible enrichment of gene expression within certain brain regions across multiple developmental time periods, using data from the Brainspan Atlas of the Developing Human Brain (58, 59). See Supplementary Methods.

## RESULTS

We performed WES on 129 CMS parent-child trios (387 samples total) meeting inclusion criteria. WES data from 853 unaffected control trios, already sequenced from the Simons Simplex Collection, were pooled with our CMS trios for joint variant calling. After quality control methods, our sample size for a burden analysis was 118 CMS and 750 unaffected trios (Table 1, Figure 1, Table S1, Figure S1).

**Table 1.** – Distribution of de novo variants in CMS cases and controls

| De novo variant type<sup>a</sup> | Variant counts |  | Mutation rate (x10<sup>-8</sup>) per bp (95% CI)<sup>j</sup> |  | Estimated coding variants per individual (95% CI)<sup>k</sup> |  | Rate ratio (95% CI) | p-value<sup>l</sup> |
| --- | --- | --- | --- | --- | --- | --- | --- | --- |
|  | CMS (N=118) | Control (N=750) | CMS (N=118) | Control (N=750) | CMS (N=118) | Control (N=750) |  |  |
| All<sup>b</sup> | 134 | 666 | 1.91<br>(1.60-2.27) | 1.68<br>(1.56-1.82) | 1.29<br>(1.08-1.54) | 1.14<br>(1.05-1.23) | 1.14<br>(0.97-1.33) | 0.10 |
| Coding<sup>c</sup> | 128 | 628 | 2.02<br>(1.68-2.40) | 1.74<br>(1.61-1.88) | 1.37<br>(1.14-1.62) | 1.18<br>(1.09-1.27) | 1.16<br>(0.98-1.36) | 0.07 |
| Synonymous SNV | 30 | 171 | 0.47<br>(0.32-0.68) | 0.47<br>(0.41-0.55) | 0.32<br>(0.22-0.46) | 0.32<br>(0.28-0.37) | 1.00<br>(0.70-1.40) | 0.50 |
| Nonsynonymous<sup>d</sup> | 96 | 446 | 1.51<br>(1.23-1.85) | 1.24<br>(1.12-1.36) | 1.02<br>(0.83-1.25) | 0.84<br>(0.76-0.92) | **1.23**<br>**(1.01-1.48)** | **0.04** |
| All Missense (Mis) | 84 | 411 | 1.32<br>(1.06-1.64) | 1.14<br>(1.03-1.25) | 0.89<br>(0.72-1.11) | 0.77<br>(0.70-0.85) | 1.16<br>(0.95-1.42) | 0.12 |
| Mis-D<sup>e</sup> | 42 | 190 | 0.66<br>(0.48-0.90) | 0.53<br>(0.45-0.61) | 0.45<br>(0.32-0.61) | 0.36<br>(0.30-0.41) | 1.26<br>(0.93-1.68) | 0.11 |
| Mis-P<sup>f</sup> | 15 | 79 | 0.24<br>(0.13-0.39) | 0.22<br>(0.17-0.27) | 0.16<br>(0.088-0.26) | 0.15<br>(0.12-0.18) | 1.08<br>(0.64-1.75) | 0.43 |
| Mis-B<sup>g</sup> | 25 | 137 | 0.39<br>(0.26-0.58) | 0.38<br>(0.32-0.45) | 0.26<br>(0.18-0.39) | 0.26<br>(0.22-0.30) | 1.04<br>(0.70-1.50) | 0.46 |
| Likely Gene Disrupting (LGD)<sup>h</sup> | 12 | 35 | 0.19<br>(0.098-0.33) | 0.097<br>(0.068-0.13) | 0.13<br>(0.066-0.22) | 0.066<br>(0.046-0.088) | **1.95**<br>**(1.04-3.50)** | **0.04** |
| Damaging (LGD + Mis-D) | 54 | 225 | 0.85<br>(0.64-1.11) | 0.62<br>(0.54-0.71) | 0.58<br>(0.43-0.75) | 0.42<br>(0.37-0.48) | **1.37**<br>**(1.05-1.76)** | **0.03** |
| LGD SNV | 10 | 19 | 0.16<br>(0.076-0.29) | 0.053<br>(0.032-0.082) | 0.11<br>(0.051-0.20) | 0.036<br>(0.022-0.055) | **3.00**<br>**(1.43-6.03)** | **0.007** |
| LGD Stopgain | 6 | 16 | 0.095<br>(0.035-0.21) | 0.044<br>(0.025-0.072) | 0.064<br>(0.024-0.14) | 0.030<br>(0.030-0.049) | 2.13<br>(0.82-5.02) | 0.10 |
| LGD Splice | 4 | 3 | 0.063<br>(0.017-0.16) | 0.0083<br>(0.0017-0.024) | 0.043<br>(0.011-0.11) | 0.0056<br>(0.0012-0.016) | **7.59**<br>**(1.66-38.5)** | **0.01** |
| LGD frameshift indel | 2 | 16 | 0.032<br>(0.0038-0.11) | 0.044<br>(0.025-0.072) | 0.022<br>(0.0026-0.074) | 0.030<br>(0.017-0.049) | 0.71<br>(0.12-2.56) | 0.77 |
| **Nonframeshift indel** | 1 | 3 | 0.016<br>(0.0004-0.088) | 0.0083<br>(0.0017-0.024) | 0.011<br>(0.00027-0.060) | 0.0056<br>(0.0012-0.016) | 1.90<br>(0.07-17.2) | 0.47 |
| **Unknown<sup>i</sup>** | 1 | 8 | 0.016<br>(0.0004-0.088) | 0.022<br>(0.010-0.041) | 0.011<br>(0.00027-0.060) | 0.015<br>(0.0068-0.028) | 0.71<br>(0.03-4.28) | 0.77 |

### Increased burden of de novo damaging variants in CMS

Based on work in other neurodevelopmental disorders, we expected to find an enrichment of de novo LGD variants (stop codon, frameshift, or canonical splice-site variants) in CMS probands versus controls. We found a statistically significant increased rate of de novo LGD variants in CMS cases, confirming our hypothesis (rate ratio [RR] 1.95, 95% Confidence Interval [CI] 1.04-3.50, p=0.04). Furthermore, de novo variants predicted to be damaging (LGD plus missense variants with Polyphen2-HDIV score <0.957 and ≥0.453) were also over-represented in CMS probands (RR 1.37, CI 1.05-1.76, p=0.03). We did not detect a difference in mutation rates for de novo synonymous variants, or when all de novo variants (coding +/- non-coding) were considered together. (Table 1, Figure 2, Table S2)

**Figure 2.**
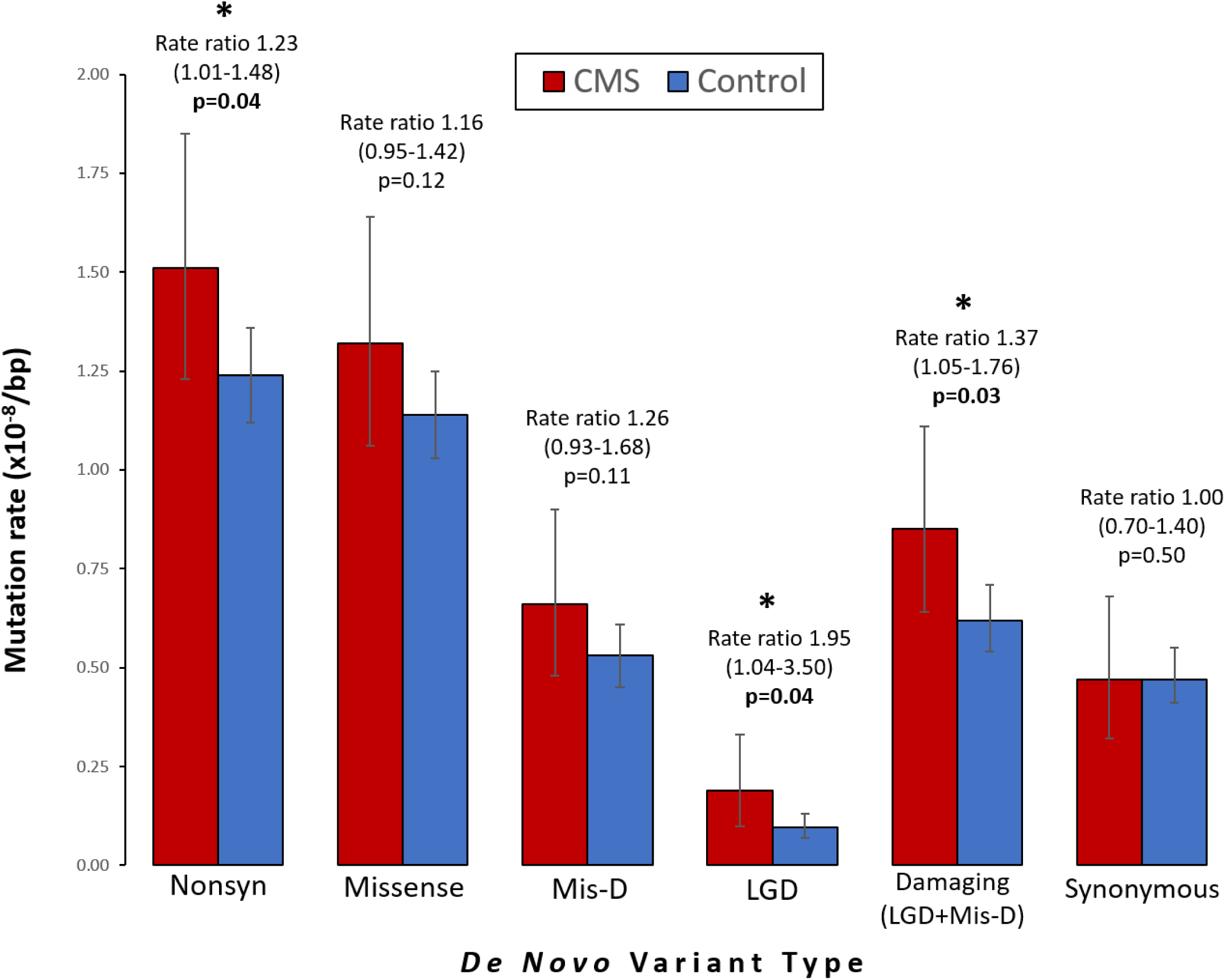
– Rates of de novo variants in CMS cases versus controls. Bar chart comparing the rates of de novo variant classes between CMS cases (red) and controls (blue). Comparisons are between per base pair (bp) mutation rates, using a one-tailed rate ratio test. Statistically significant comparisons (p<0.05) are marked with asterisks. Error bars show 95% confidence intervals.

### *KDM5B* is a high-confidence candidate risk gene in CMS

Having established a higher rate of de novo damaging variants in CMS probands, we next asked whether these variants cluster within specific genes. We identified one gene with more than one predicted damaging de novo variant in unrelated probands: *KDM5B* (*Lysine Demethylase 5B)* harbored two different LGD (stopgain) de novo variants in CMS probands 1029-03 and 1050-03. Using TADA-Denovo (53) and previously established false discovery rate (FDR) thresholds, we found that *KDM5B* meets statistical criteria for a high-confidence risk gene (q<0.1) in CMS (Table S3).

### Approximately 184 genes contribute to CMS risk

Based on the number of observed de novo damaging mutations in CMS, the MLE method estimated the most likely number of CMS risk genes to be 184 (Figure S2). Next, we used this estimate along with de novo mutation rates observed in CMS trios to predict the likely number of risk genes that will be discovered in larger CMS cohorts. Based on these simulations, WES of 500 trios should find 16 probable and 7 high-confidence risk genes; 1000 trios should find 51 probable and 26 high-confidence risk genes (Figure S3).

### CMS gene enrichment in Tourette syndrome, “suggestive evidence” ASD genes, and in ASD with high stereotypy scores

Using DNENRICH (56), we found significant overlap between genes harboring de novo damaging variants in CMS (52 genes after excluding two genes with de novo damaging variants in controls) and several gene sets curated from the literature (Table S4). In particular, our CMS cohort genes show significant gene overlap with autism probands with high stereotypy scores (5.8x enrichment, p=0.048), Tourette’s disorder (4.7x enrichment, p=0.011), and category 3 (suggestive evidence) autism risk genes (3.7x enrichment, p=0.023). There was no significant overlap with OCD, ADHD, schizophrenia, intellectual disability, developmental disabilities, or other categories of autism gene lists (Table S4).

### Exploratory pathway, gene ontology, and spatiotemporal analyses

Using this same list of 52 genes harboring de novo damaging variants in CMS, we performed exploratory analyses to determine shared underlying canonical pathways and functional connectivity. Our input gene list is significantly enriched for biological process ontology-based sets including demethylation, organization of branching structures (e.g. neurons and vasculature), multicellular organism development, and cell motility. Enriched canonical pathways include the glucocorticoid receptor pathway, transient receptor potential (TRP) channel function, demethylation of histones by histone lysine demethylases, and ion channel transport (Table S5).

Finally, mapping our CMS de novo damaging variant genes onto the Brainspan Atlas of the Developing Human Brain gene expression data, we see nominal enrichment in early mid-fetal cortex and striatum, with a possible trends toward enrichment in early fetal hippocampus, late mid-fetal cerebellum, and young childhood cerebellum (Table S5).

## DISCUSSION

Like prior studies of ASD, Tourette’s disorder, and OCD, the current study demonstrates that the identification of de novo variants will identify risk genes and provide a reliable entry-point into understanding the biology of stereotypies. We are studying otherwise typically-developing children with stereotypies (primary CMS), as this may represent a more genetically homogenous group of individuals versus those with secondary stereotypies, thereby facilitating genetic discovery and insight into the biology of stereotypies more generally (60, 61). Despite our small cohort size, we identified two de novo nonsense mutations in *KDM5B* in unrelated probands, and we show that finding two such mutations in our cohort is highly unlikely to be a chance occurrence.

*KDM5B* is a lysine-specific demethylase that removes methyl groups from tri-, di- and monomethylated lysine 4 on histone 3. *KDM5B* acts a transcriptional repressor and has primarily been implicated in the pathogenesis of cancer (62). More recently, this gene has also been implicated in congenital heart disease risk, embryonic stem cell renewal and differentiation, and DNA repair (63–65). *KDM5B* has been identified as high-confidence risk gene in WES studies of ASD (54), and expression is normally restricted to the brain and the testis (66). Within the brain, high expression levels are in the cerebellum (Figure S4), and expression across all brain regions is highest prenatally (Figure S5). The identification of this risk gene in CMS suggests that chromatin (dys)regulation of *KDM5B* target genes may be one contributing mechanism underlying stereotypies. As shown in our gene ontology and pathway results (Table S5), demethylation of histones is enriched, driven by the two *KDM5B* mutations and a de novo damaging mutation in *KDM3B* in a third proband. Certainly, further studies are warranted to determine the downstream effects of these mutations in the developing brain. These studies are underway in our laboratory.

It is interesting that we find significant overlap between genes harboring de novo damaging mutations in CMS and those reported in a recent study of Tourette syndrome (Table S4; 4.7x enrichment, p=0.011). We find this overlap with Tourette despite excluding CMS subjects with comorbid motor or vocal tics (see Methods). Enriched expression of CMS genes in the cortex and striatum (Table S5) is also consistent with widely believed involvement of these regions in Tourette syndrome. While OCD and ADHD symptoms were not exclusionary in our CMS study, we saw no gene overlap with these disorders. Similarly, we found no overlap with SCZ, ID, or DD. We did, however, find significant overlap between CMS and Category 3 (suggestive evidence) ASD risk genes (3.7x enrichment, p=0.023).

With regard to stereotypies in ASD, we curated lists of genes harboring de novo damaging mutations in SSC probands with the highest (90^th^ percentile) and lowest (10^th^ percentile) stereotypies, measured by Stereotyped Behavior Scores (SBS) from the RBS-R. *KDM5B* mutations were found only in SSC probands with high stereotypy scores, yielding 5.8-fold enrichment over expectation (p=0.047) when compared against our CMS genes (Table S4). To further examine the relation between de novo *KDM5B* mutations stereotypies in SSC ASD probands, we compared SBS scores in four probands with *KDM5B* mutations versus 364 age-matched patients without (Figure S6). Scores were higher in mutation carriers, but this did not reach statistical significance (p=0.076), likely due to the low number of mutation carriers in this cohort.

In summary, we report an increased burden of de novo damaging DNA coding variants in primary complex motor stereotypies. We identified one high-confidence risk gene for CMS in our pilot cohort and estimate that there are 184 genes conferring risk for this phenotype. Whole-exome sequencing in parent-child CMS trios provides a reliable way to make progress in gene discovery. Our preliminary analyses of genes harboring de novo damaging mutations in CMS highlight several biological pathways, processes, brain regions, and developmental time periods that give insights into possible etiologies of stereotypies, which are a prerequisite to development of new treatments. Further sequencing and mechanistic studies are warranted to understand this phenotype, which has relevance across diagnostic boundaries.

## ACKNOWLEDGEMENTS

We wish to thank the families who have participated in and contributed to this study. Control subject data were obtained from the NIH-supported National Database for Autism Research (NDAR). NDAR is a collaborative informatics system created by the National Institutes of Health to provide a national resource to support and accelerate research in autism. Dataset identifier: 2042. This manuscript reflects the views of the authors and may not reflect the opinions or views of the NIH or of the Submitters submitting original data to NDAR. This work was supported by grants from the Simons Foundation (SFARI award #239013, TVF), the Allison Family Foundation (TVF), Nesbitt-McMaster Foundation (HSS), Klump Family (HSS), and Graves Family (HSS). We are grateful to all of the families at the participating Simons Simplex Collection (SSC) sites, as well as the principal investigators (A. Beaudet, R. Bernier, J. Constantino, E. Cook, E. Fombonne, D. Geschwind, R. Goin-Kochel, E. Hanson, D. Grice, A. Klin, D. Ledbetter, C. Lord, C. Martin, D. Martin, R. Maxim, J. Miles, O. Ousley, K. Pelphrey, B. Peterson, J. Piggot, C. Saulnier, M. State, W. Stone, J. Sutcliffe, C. Walsh, Z. Warren, E. Wijsman). We appreciate obtaining access to phenotype and genetic data on SFARI Base. Approved researchers can obtain the SSC population dataset described in this study (https://www.sfari.org/resource/simons-simplex-collection/) by applying at https://base.sfari.org.

Dr. Fernandez receives research/grant support from the National Institutes of Mental Health and the Brain and Behavior Research Foundation. Dr. Olfson receives research support from the American Academy of Child & Adolescent Psychiatry and the Alan B. Slifka Foundation through the Riva Ariella Ritvo endowment. Dr Singer serves as a consultant for Abide Therapeutics, Inc; Cello Health BioConsulting; ClearView Healthcare Partners; Teva Pharmaceutical Industries Ltd; and Trinity Partners, LLC. Dr Singer receives publishing royalties from Elsevier and research/grant support from the Tourette Association of America. Other authors declare no potential conflicts.

## SUPPLEMENTARY METHODS

### Subjects and Assessment Measures

The Stereotypy Severity Scale (SSS) is a 5-item caregiver questionnaire consisting of two components (Motor and Impairment) for the ranking of motor stereotypy severity (1). The SSS Motor score (range: 0-18) quantifies motor severity and rates movements along four discriminate dimensions: number (0–3), frequency (0–5), intensity (0–5), and interference (0–5). The SSS Impairment score (range 0-50) is an independent rating of difficulties in self-esteem, family, school, or social acceptance caused by the movements.

The ASSQ is a 27-item caregiver questionnaire addressing symptoms of ASD (2).

Other parent-completed measures included: the Multidimensional Anxiety Scale for Children (MASC), assessing symptoms of anxiety (3); the ADHD-Rating Scale-IV (4) and Conners Parent Rating Scale (CPRS), assessing symptoms of ADHD; the Child Yale Brown Obsessive-Compulsive Scale (CY-BOCS), assessing symptoms of OCD (5); the Repetitive Behavior Scale-Revised (RBS-R), assessing repetitive behaviors (6); and the Social Responsiveness Scale (SRS), assessing social communication skills (7).

### Whole-exome sequencing, alignment, variant calling, and quality control

Exome capture and sequencing were performed at the Yale Center for Genome Analysis (YCGA), using the NimbleGen SeqCap EZExomeV2 capture library (Roche NimbleGen, Madison, WI, USA) and the Illumina HiSeq 2500 platform (74 bp paired-end reads; Illumina, San Diego, CA, USA). We multiplexed six samples during each capture reaction and sequencing lane, pooling parents and probands when possible. Alignment and variant calling of the sequencing reads followed the latest Genome Analysis Toolkit (GATK) (8) Best Practices guidelines, as described previously (9). Reads were aligned using BWA-mem (10) to the b37 human reference sequence with decoy sequences. Picard’s MarkDuplicates tool was used to mark PCR duplicates (https://broadinstitute.github.io/picard/). GATK was used to realign indels, recalibrate quality scores, and generate GVCF files for each sample using the HaplotypeCaller tool. All samples were called jointly using GATK’s GenotypeGVCFs tool, variant score recalibration was applied to the called variants, and all variant call data was written to a VCF file. This pipeline uses GATK’s Best Practices parameters and the default parameters for BWA and Picard. Only passing variants were used in downstream analyses. Variants were annotated using the RefSeq hg19 gene definitions and multiple external databases of variant population frequency, conservation scores, variation intolerance, mutation severity, and predicted functional effects using ANNOVAR (11).

Relatedness statistics were calculated based on the method of Manichaikul et al. (12), implemented in VCFtools v0.1.14.10 (13). Trios were omitted if expected family relationships were not confirmed or if there were unexpected relationships within or between families. Trios were omitted if > 5 de novo variants were observed. PLINK/SEQ (14) (i-stats; https://psychgen.u.hpc.mssm.edu/plinkseq/stats.shtml), PicardTools, and GATK DepthOfCoverage tools were used to generate quality metrics (Table S1). To identify outliers that might confound our case-control analysis, we performed principal components analysis (PCA) using this data. A scree plot determined the number of principal components accounting for the greatest proportion of variance, and we removed trios with family members falling more than three standard deviations from the mean in any of these principal components (Figure S1, Table S1). The R code PCA is provided below.

We used stringent thresholds for identifying de novo mutations because DNA from control subjects in the Simons Simplex Collection was not available for confirmation by Sanger sequencing. As previously described (15), de novo variants were called using an in-house script that required: (a) child is heterozygous for a variant, with alternate allele frequency between 0.3 and 0.7 in the child and < 0.05 in the parents; (b) sequencing depth (DP) ≥ 20 in all family members at the variant position; (c) alternate allele depth (AD) ≥ 5; (d) observed allele frequency (AC) < 0.01 (1%) among all cases and controls; and (e) mapping quality (MQ) ≥ 30. False positive calls were removed by in silico visualization. Based on Sanger sequencing confirmations, our confirmation rate using this method and platform is 98.7% (16).

### Principal Component Analysis (PCA)

PCA was performed on all sequencing quality metrics (Table S1) in R using the following code:

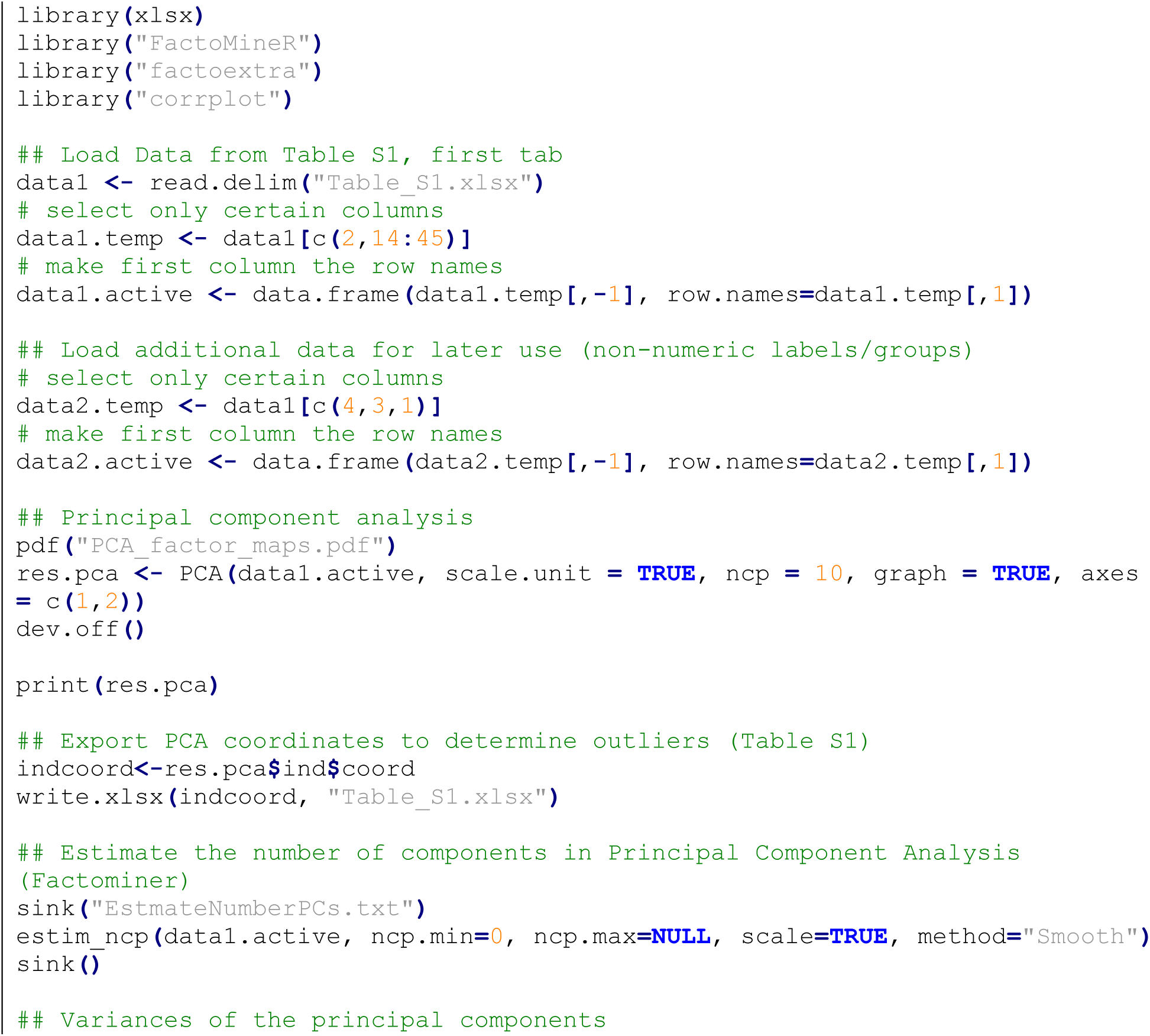

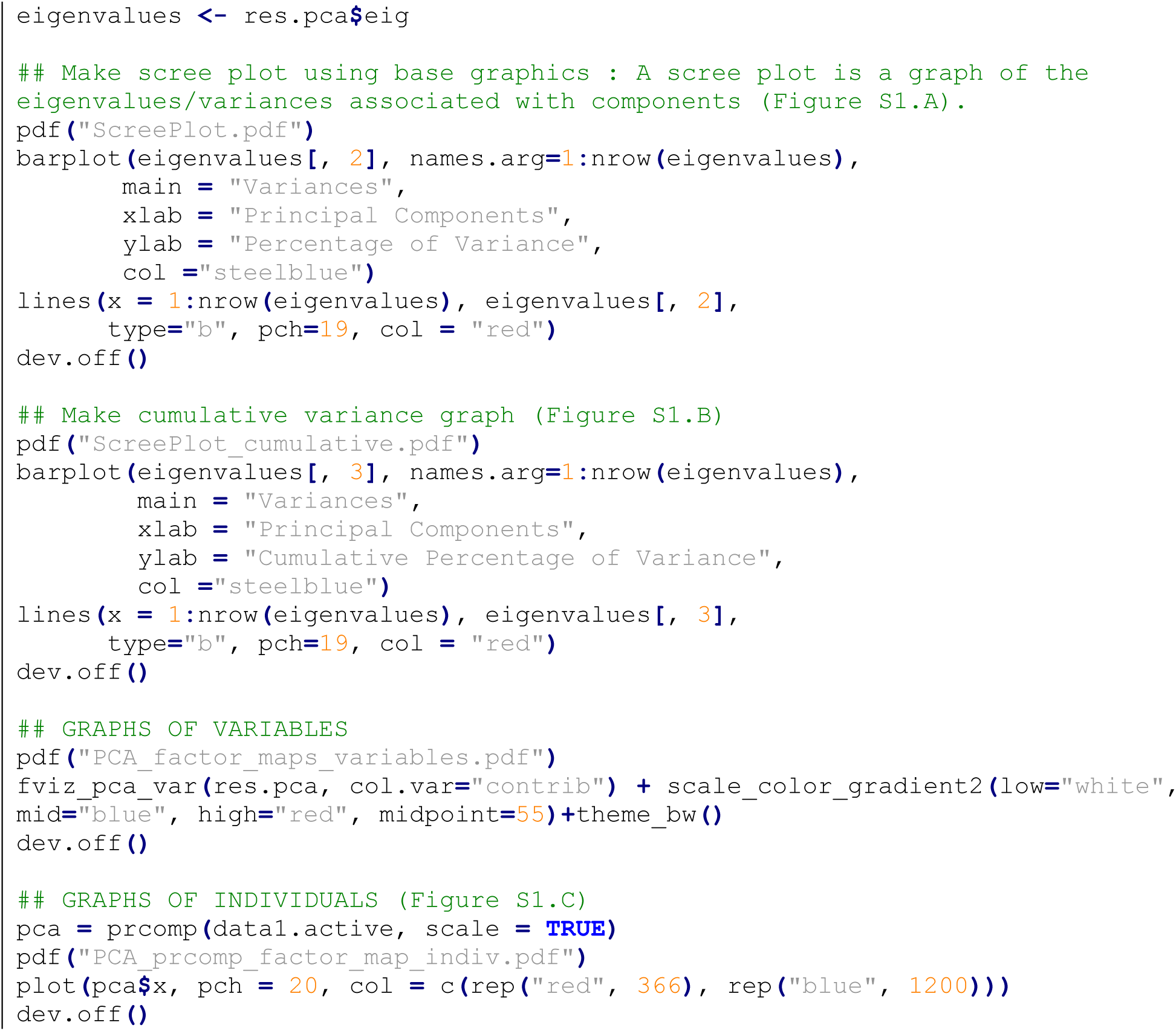

### Mutation rate calculations

Within each cohort, we calculated the rate of de novo mutations per base pair. For accurate rate calculation, we first determined the number of “callable” base pairs per family using the GATK DepthOfCoverage tool. We considered only bases covered at ≥ 20x in all family members, with base quality ≥ 20, and map quality ≥ 30; these thresholds match those required for GATK and de novo variant calling. For each cohort, we summed the “callable” base pairs in every family and used this number as the denominator for de novo rate calculations. Details for calculating these callable base pairs is below. The resulting rate was divided by two to give haploid rates. Confidence intervals were calculated using the *pois.conf.int (pois.exact)* function from the epitools v0.5-9 package in R. We compared de novo mutation rates in cases versus controls (burden analysis) using a one-tailed rate ratio test in R (https://cran.r-project.org/package=rateratio.test), considering only those variants present with a frequency of <0.001 in the ExAC v0.3.1 database (17).

### Calculating “callable” base pairs

The following command was used to calculate the callable base pairs in each trio:

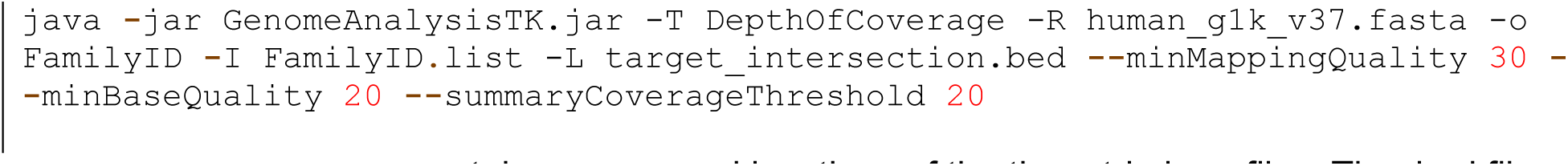

`FamilyID.list` contains names and locations of the three trio bam files. The .bed file contains the genomic intervals over which to calculate the callable base pairs. To calculate the coding callable base pairs (used for coding mutation rates, e.g. synonymous, nonsynonymous, missense, etc., see Table 1, Table S2), we used a bed file with intervals spanning the intersection of both capture array target intervals and the RefSeq coding intervals (32,027,823 bp total). To calculate all callable base pairs (used for the total coding + noncoding mutation rate, see “All” in Table 1), we used a bed file with intervals spanning the intersection of both capture array target intervals (33,973,867 bp total). The number of coding and total callable base pairs for every family passing is listed in Table S1.

### Calculating expected mutation rates for downstream analyses

To perform subsequent maximum likelihood estimation (MLE) and TADA analyses, we used published per gene de novo mutation rates from unaffected parent-child trios (18). From the control samples in our dataset, we calculated the proportion of the overall coding mutation rate that comprised LGD and Mis-D mutations, and then used these proportions to calculate the expected LGD and Mis-D mutation rate per gene (Table S3).

The following R code was used to generate the expected mutation rates:

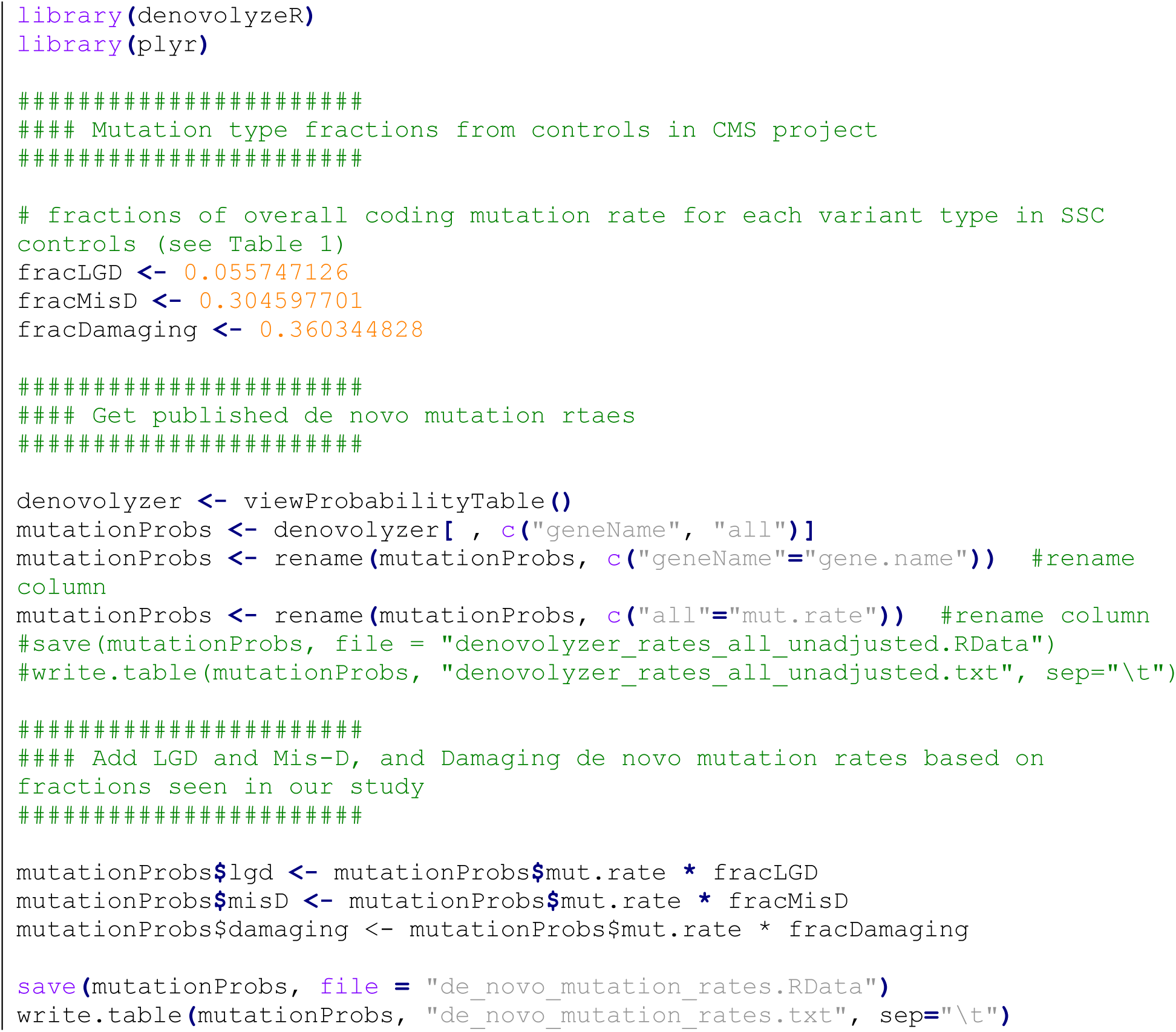

### TADA analysis

As shown in prior WES studies in neuropsychiatric disorders, a small number of rare de novo mutations in the same gene among unrelated individuals can provide considerable statistical power to establish association. To test this hypothesis in CMS, we used the transmitted and de novo association (TADA-Denovo) test. TADA uses a Bayesian model that combines data from de novo mutations and population mutation rates to increase the power of gene discovery. While TADA has a version that can include inherited variants, we did not include inherited data in this study, because their confirmation rate is not known and their contribution to the TADA score is minimal, given their lower relative risks (19, 20). The code and documentation for this tool can be found here (http://wpicr.wpic.pitt.edu/WPICCompGen/TADA/TADA_homepage.htm).

We describe the parameters of this test in our prior WES studies of Tourette’s disorder and OCD (15, 21). The code and parameters used in the current study are given below. A low FDR-corrected q-value represents strong evidence for association. Genes with FDR q<0.3 are considered probable risk genes, and those with FDR q<0.1 are high-confidence risk genes.

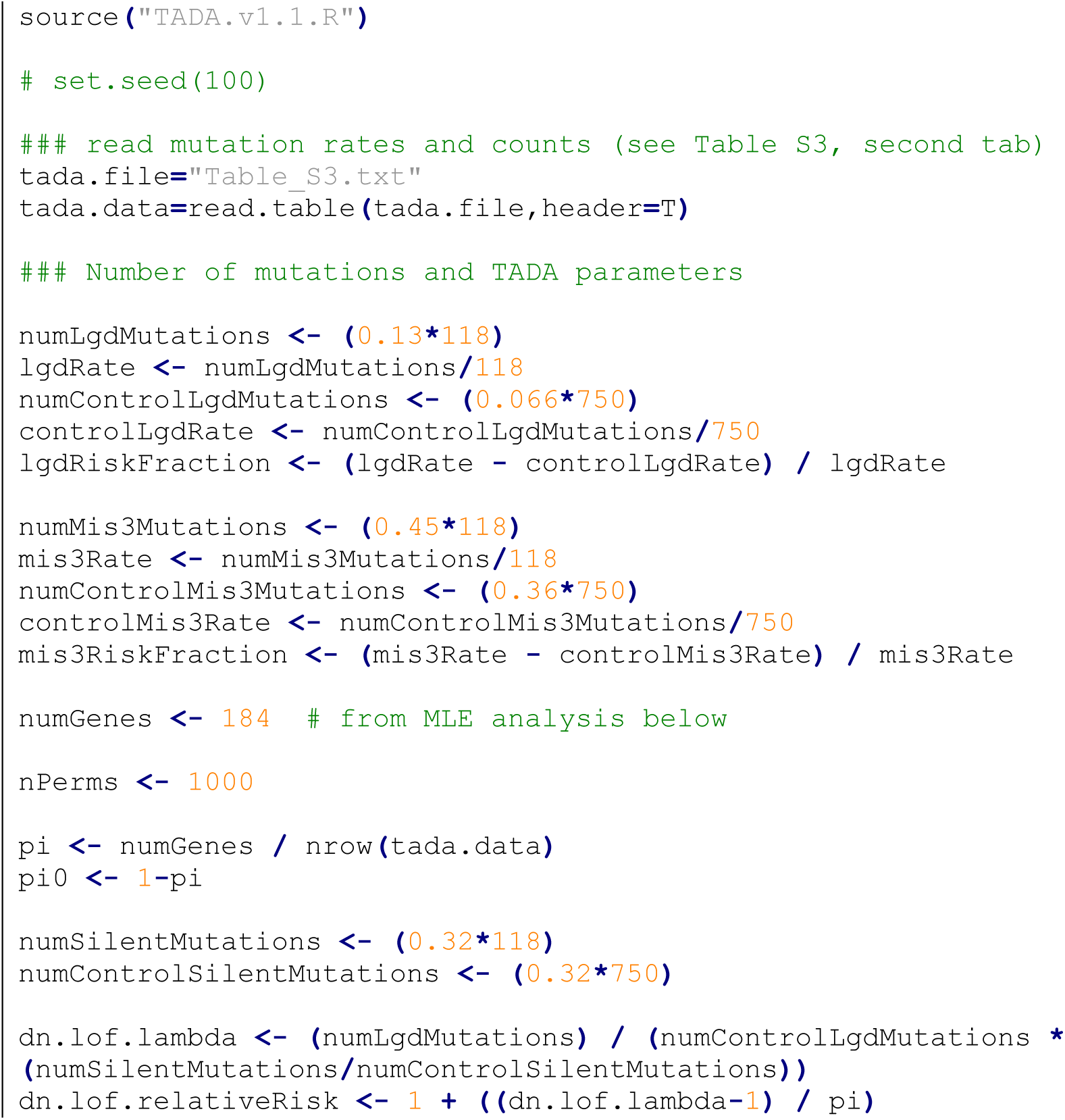
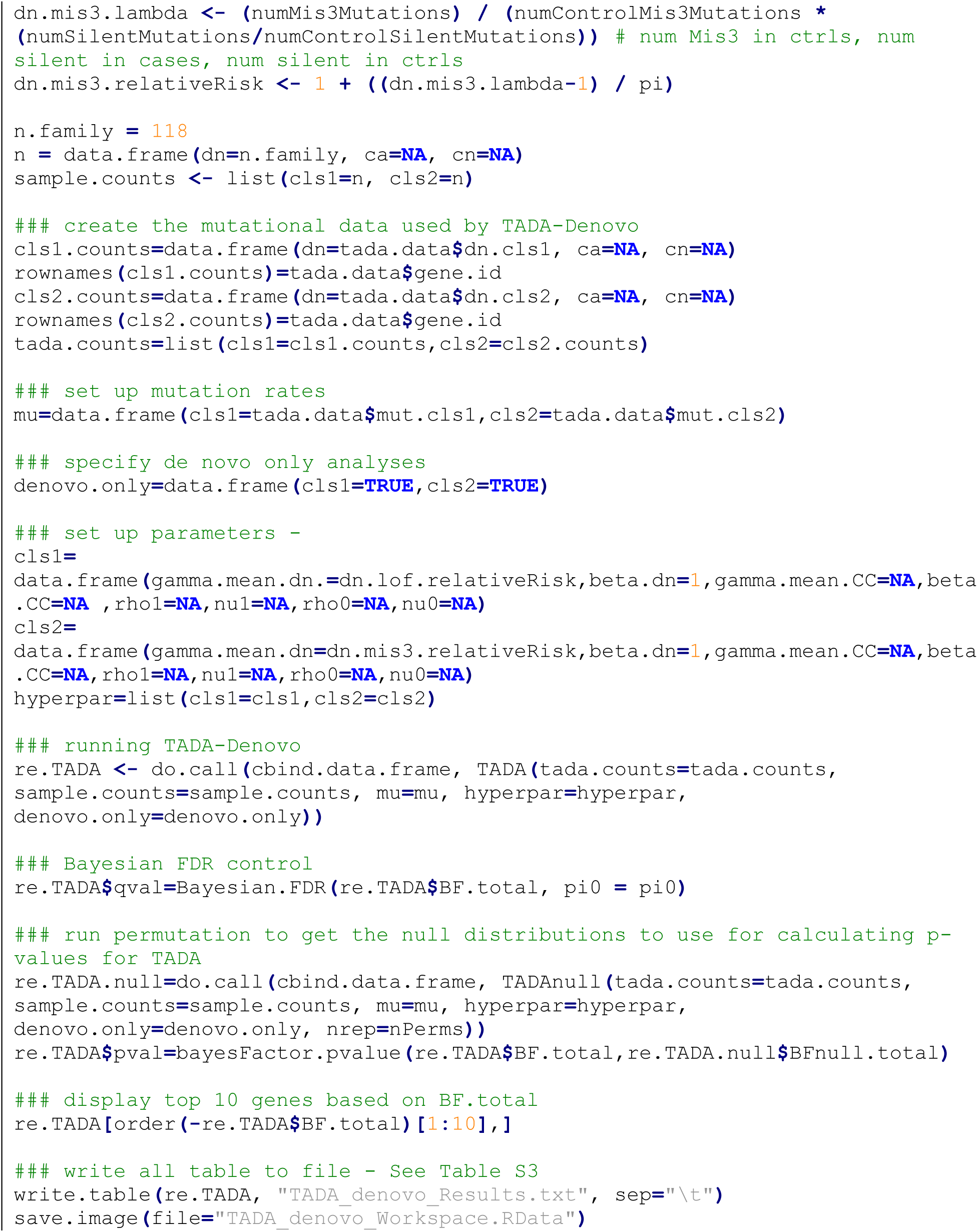

### Maximum Likelihood Estimation (MLE) method for estimating the number of CMS risk genes

We used a maximum likelihood estimation (MLE) method to estimate the number of genes contributing risk to CMS, based on vulnerability to de novo damaging variants (22). For every number of risk genes from 1 to 2,500, we simulated 54 variants (the number of damaging de novo variants observed in probands in our case-control burden analysis). Variant simulations were performed 50,000 times at each number of risk genes. Following each simulation, a percentage of variants was randomly assigned to the risk genes. The percentage of variants assigned to risk genes was determined by the fraction of de novo damaging variants estimated to carry CMS risk, and variant simulations were weighted by gene size and GC content (19). We then counted the number of risk and non-risk genes containing two variants and the number containing three or more variants. The frequency of concordance between our simulated and observed data was calculated. A curve was plotted to show the concordance frequency (y-axis) at each assumed number of risk genes (x-axis), and the peak was taken as the estimate of the most likely number of risk genes (22). See Figure S2.

The following R code was used to perform these calculations:

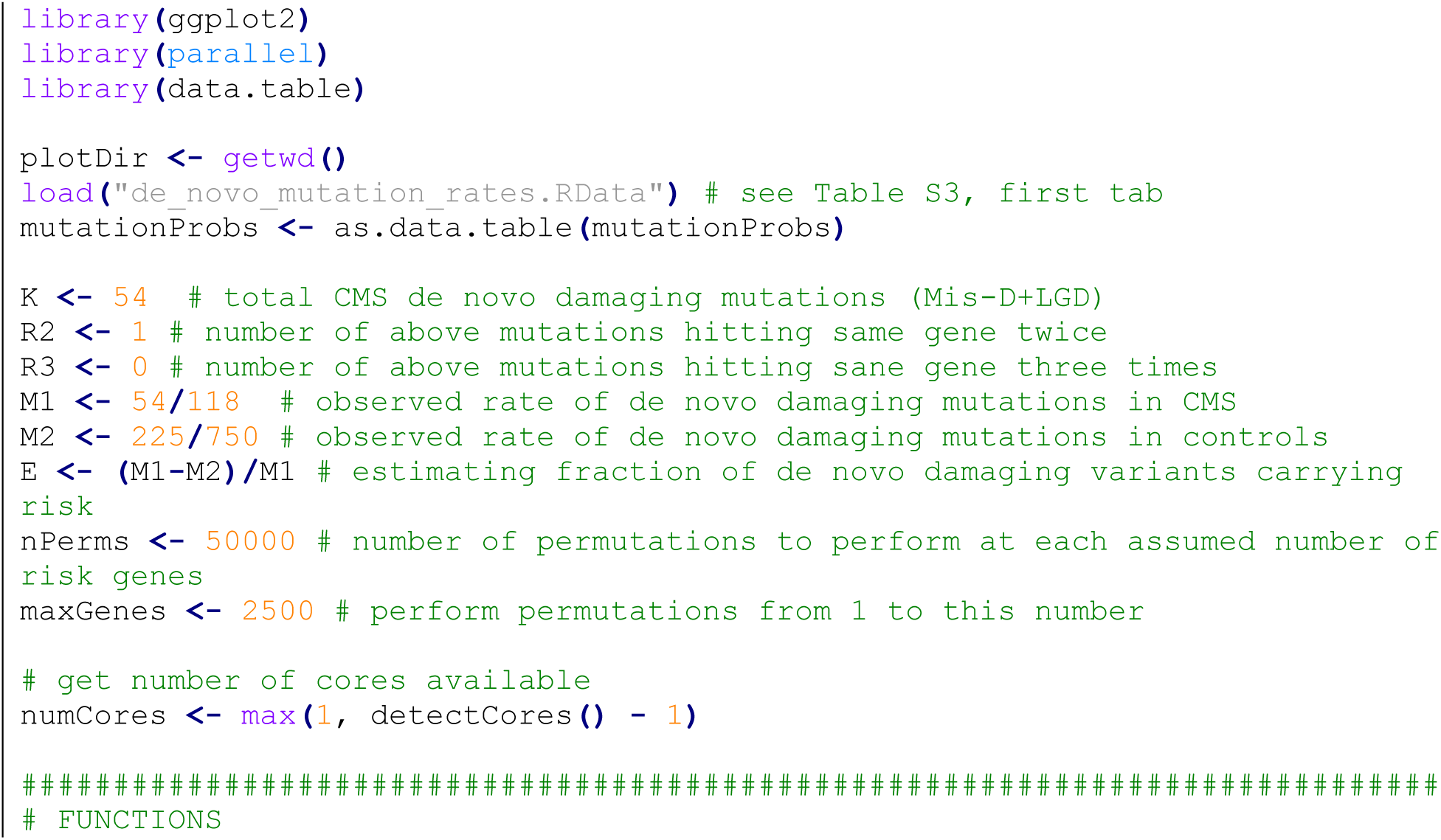
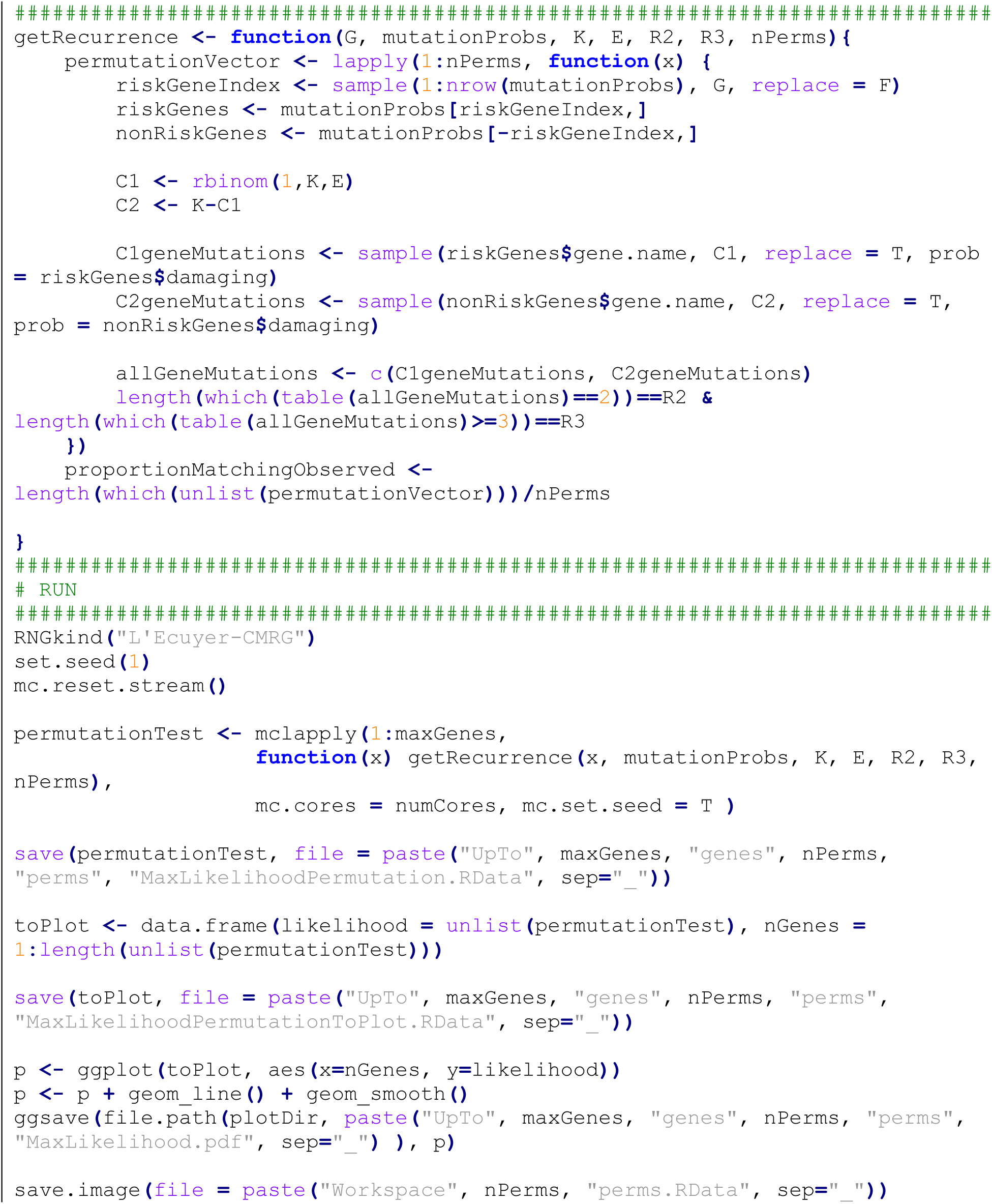

### Predicting the number of CMS risk genes identified by cohort size

Future discovery of risk genes was predicted as previously described (16, 21). Fixing the number of CMS risk genes at 184 (from estimate above), we simulated de novo mutations, with the number of mutations matching the observed mutation rate in CMS probands, and with mutation simulations weighted by gene size and GC content. Simulations were performed at each cohort size, from 25 to 3000, in increments of 25. Simulated variants were randomly assigned to the risk genes, with the percentage of variants assigned to risk genes determined by the fraction of de novo damaging variants estimated to carry CMS risk. At each cohort size, 10,000 simulations were performed. LGD and Mis-D variants were simulated separately. Simulated variants were then combined and given as input to the TADA-Denovo algorithm, using the same parameters described above for the observed data. The number of high confidence (q<0.1) and probable (q<0.3) risk genes were recorded and plotted using polynomial regression fitting; this regression model allows prediction of the number of genes identified at a specified cohort size. See Figure S3.

The following R code was used to perform these calculations:

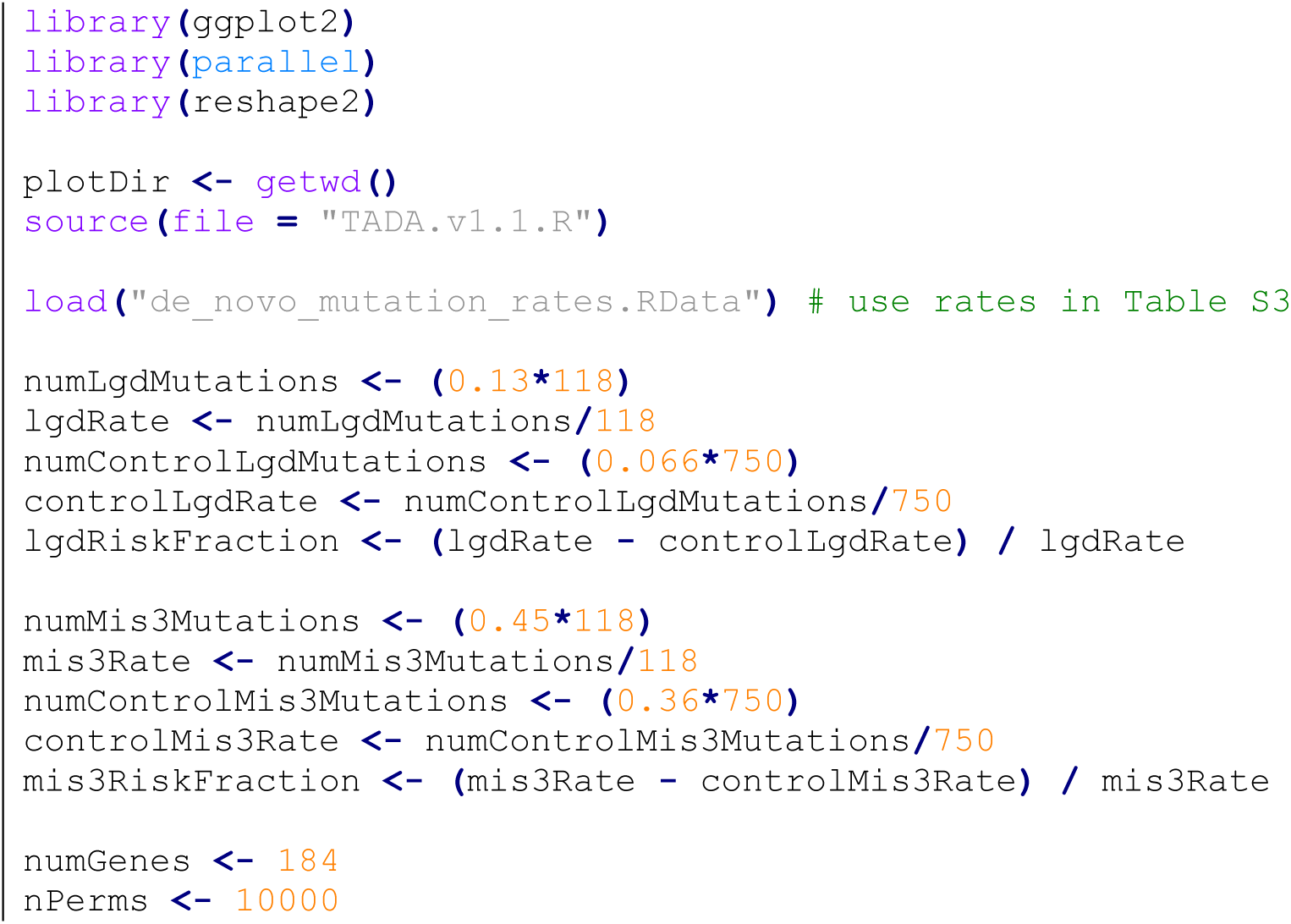
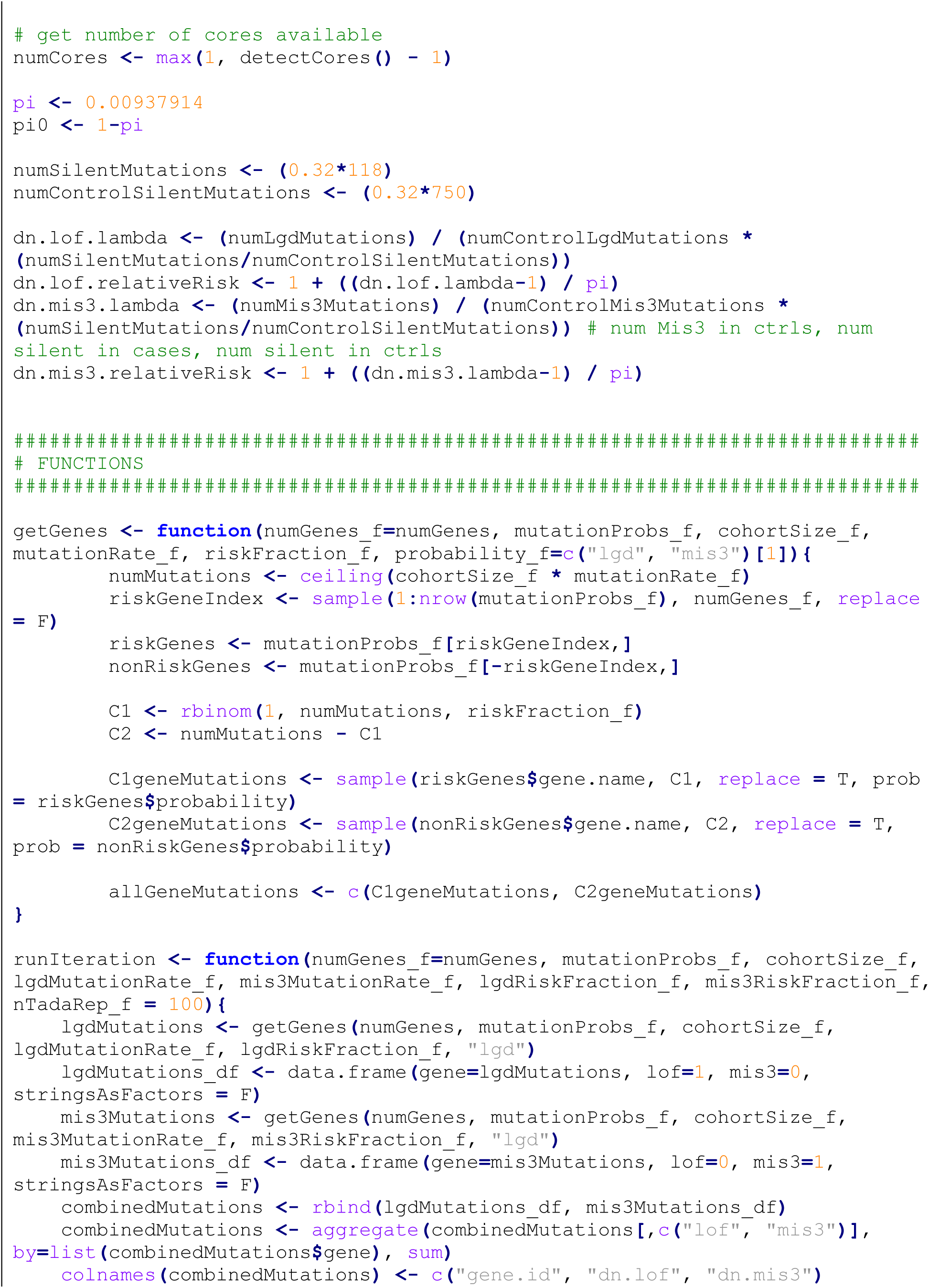
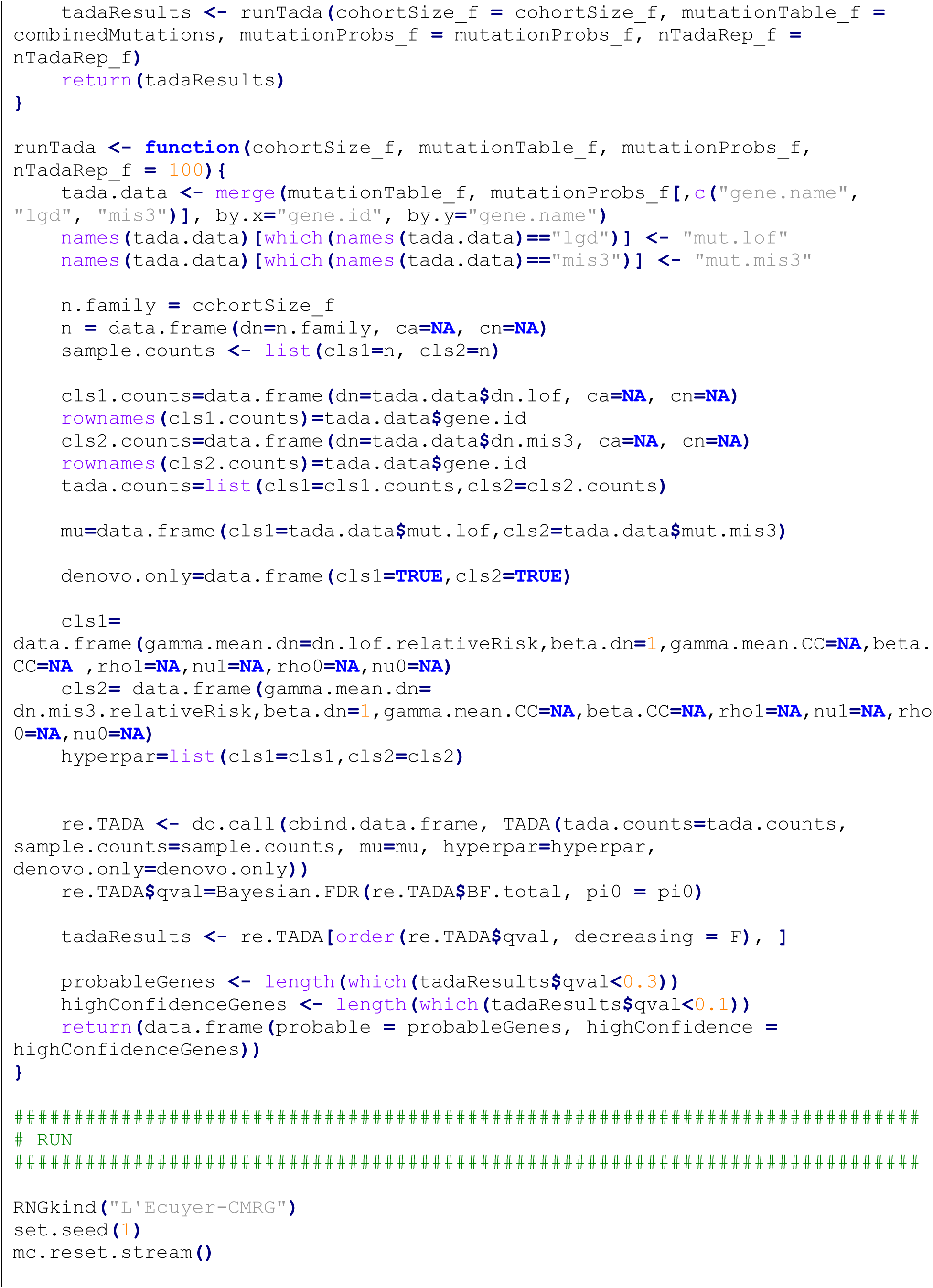
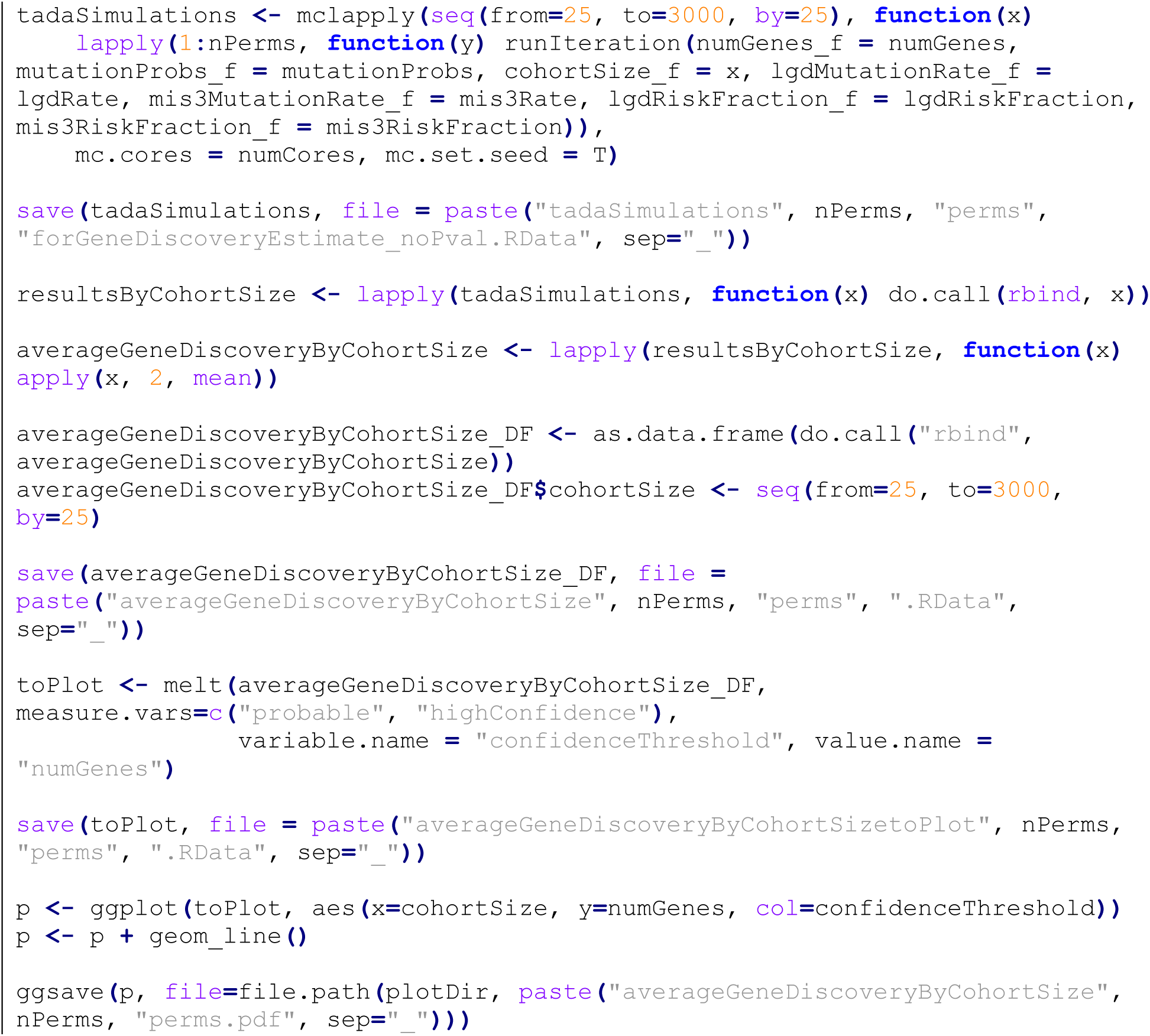

### Gene set overlap

We used DNENRICH (14) (https://psychgen.u.hpc.mssm.edu/dnenrich/) to test whether CMS genes found to have de novo damaging mutations in our study (52 genes after excluding two genes with de novo damaging variants in controls) were significantly enriched among previously reported genes in autism (ASD), schizophrenia (SCZ), developmental disorders (DD), Tourette’s disorder (TD), obsessive-compulsive disorder (OCD), attention-deficit/hyperactivity disorder (ADHD), and intellectual disability (ID). Gene lists for SCZ and ID were obtained from a recent cross-disorder study (23) that included de novo single nucleotide and indel variants from multiple exome sequencing studies (14, 24–31). DD (32), TD (33), OCD (15), and ADHD (34) genes were obtained from recently published WES studies. For the TD gene list, we removed variants reported in subjects with comorbid OCD, as this phenotype information was readily available. ASD gene lists were obtained from the SFARI Gene online database (https://gene.sfari.org/about-gene-scoring/criteria/), version 08-06-2019. ASD genes in this database have been stratified into six categories, based on manually curated strength of evidence from human genetics studies. As described in more detail on the above website, gene categories are as follows: Category 1: high confidence; Category 2: strong candidate; Category 3: suggestive evidence; Category 4: minimal evidence; Category 5: hypothesized but untested; Category 6: evidence does not support a role. Finally, we curated lists of genes harboring damaging de novo mutations in ASD probands from the Simons Simplex Collection (SSC) for whom stereotyped behavior scores (Stereotyped Behavior Score from the RBS-R, Repetitive Behavior Scale-Revised) were available. De novo mutation data from SSC probands was obtained from denovo-db (http://denovo-db.gs.washington.edu/, version 1.6.1, accessed 5/14/2019). RBS-R Stereotyped Behavior Score data was obtained from the Simons Foundation. Because we were particularly interested in the question of whether our CMS cohort share genes harboring de novo damaging mutations with SSC probands having high stereotypy scores, we assembled gene lists from SSC subjects with stereotypy scores in the 90^th^ percentile (high stereotypies) and those in the 10^th^ percentile (low stereotypies). Gene lists are provided in Table S4, first tab.

DNENRICH simulates random mutations while accounting for gene size, trinucleotide context, and mutational effect. We performed 100,000 permutations, comparing the observed and expected overlap with each gene set. Empirical p-values were generated, based on a one-sided enrichment analysis under a binomial model of greater than expected hits per gene set. We tested for overlap between our CMS genes and those in each of the mentioned gene lists. Results are provided in Table S4, third tab.

The following Linux commands were used to run the DNENRICH analysis:

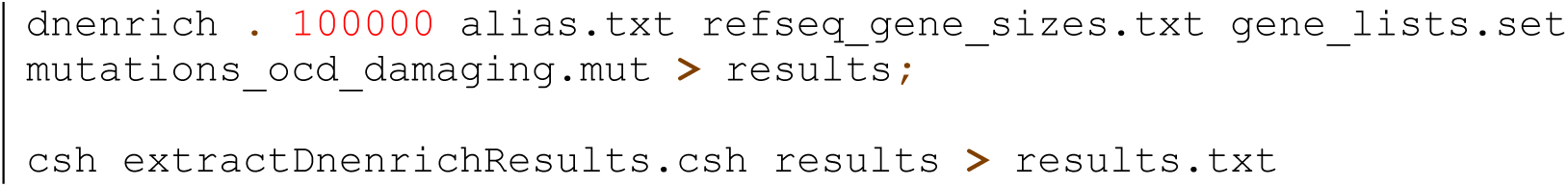

### Exploratory pathway, gene ontology, and spatiotemporal analyses

To explore whether genes harboring de novo damaging variants in our CMS probands (52 genes after excluding two genes with de novo damaging variants also found in SSC controls), we used ConsensusPathDB (35) (http://cpdb.molgen.mpg.de/, Release 34 [15.01.2019], accessed 8/9/2019). This tool integrates human protein and genetic interaction networks from 32 databases and interactions curated from the literature. The following default settings were used for ConsensusPathDB: gene set analysis → over-representation analysis; gene identifier type: gene symbol (HGNC symbol); Pathway-based sets: pathways as defined by pathway databases, select all resources, minimum overlap with input list = 2, p-value cutoff = 0.05; Gene ontology categories: gene ontology level 2 categories, select all (biological processes, molecular function, cellular component), p-value cutoff = 0.05. Results are in Table S5, first tab.

For spatiotemporal enrichment analysis, we used our same list of 52 genes and asked whether these genes have known expression patterns that cluster within certain anatomical brain regions or within certain developmental time periods. To perform this analysis, we used data from the Brainspan Atlas of the Developing Human Brain (36) as implemented in the Specific Enrichment Analysis (SEA) tool (http://genetics.wustl.edu/jdlab/csea-tool-2/, version 1.1, accessed 8/10/2019). For this analysis, we used a specificity index threshold (pSI) of 0.05 (37). Results are in Table S5, second tab. Fisher’s Exact p-values are uncorrected for multiple comparisons.

## SUPPLEMENTARY FIGURES

**Figure S1.**
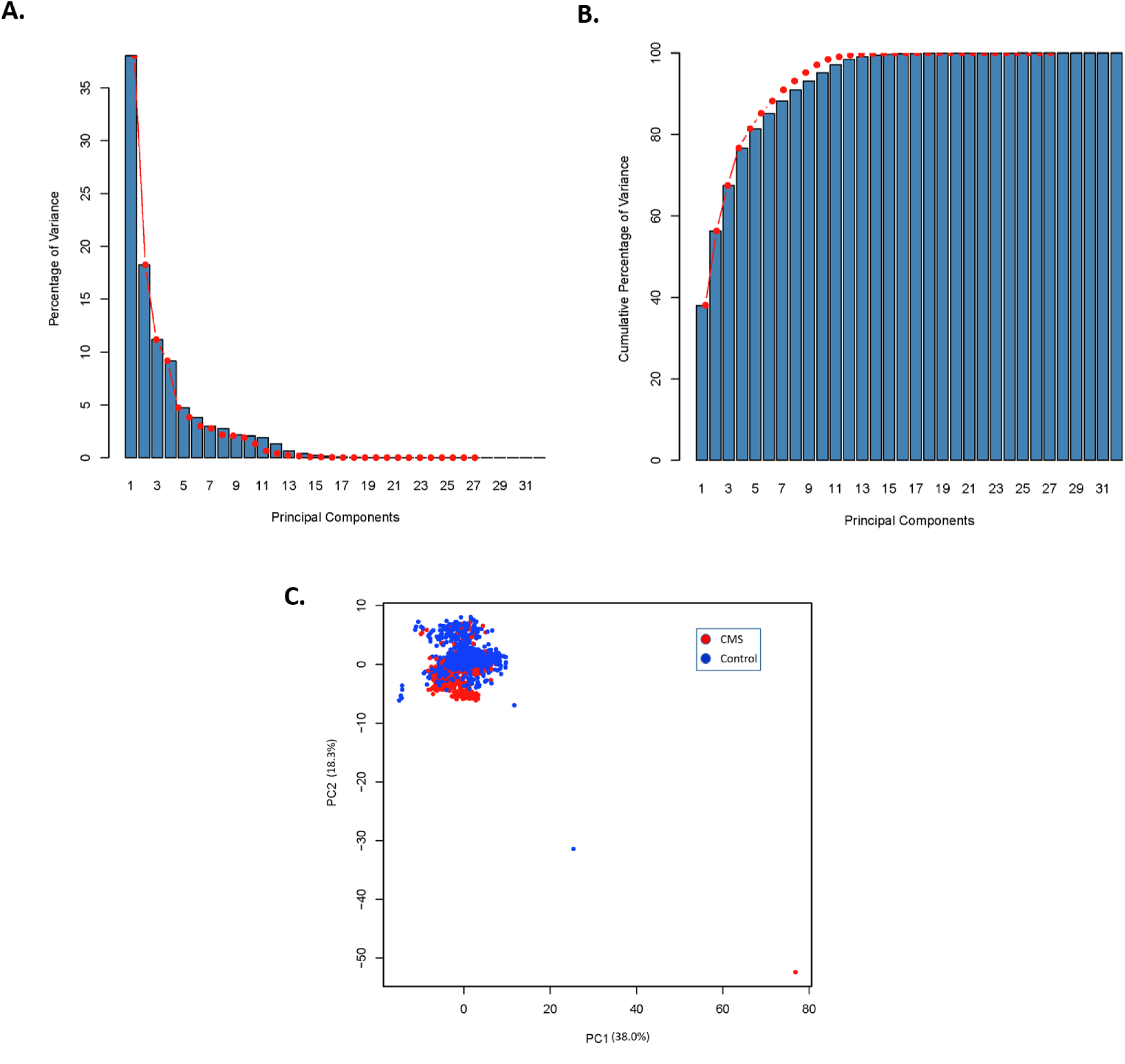
PCA scree and individual plots. Scree plots following Principal Components Analysis (PCA), showing (A) the percentage of variance captured by each of the first 32 principal components, and (B) the cumulative percentage of variance captured by these same components in the exome metrics data from cases and controls. The “elbow” of the scree plot is visualized to be around the 5^th^ principal component. This was confirmed by the Factominer R code function “estim_ncp()”. The first 5 PCs capture over 80% of the variance, and this number of PCs was used to determine PCA outliers during quality control (see Table S1 and Supplementary Methods). (C) Individual plots for the first two principal components, based on PCA of exome sequencing quality metrics. CMS cases are plotted in red, and controls in blue. The first two PCs together capture 56.3% of the variance. R code to generate this data and figure are in Supplementary Methods, and individual PC factor values are in Table S1. This figure includes PCA outliers (>3 standard deviations from the mean in PCs 1-5), which were removed during quality control, prior to further analysis of case-control data.

**Figure S2.**
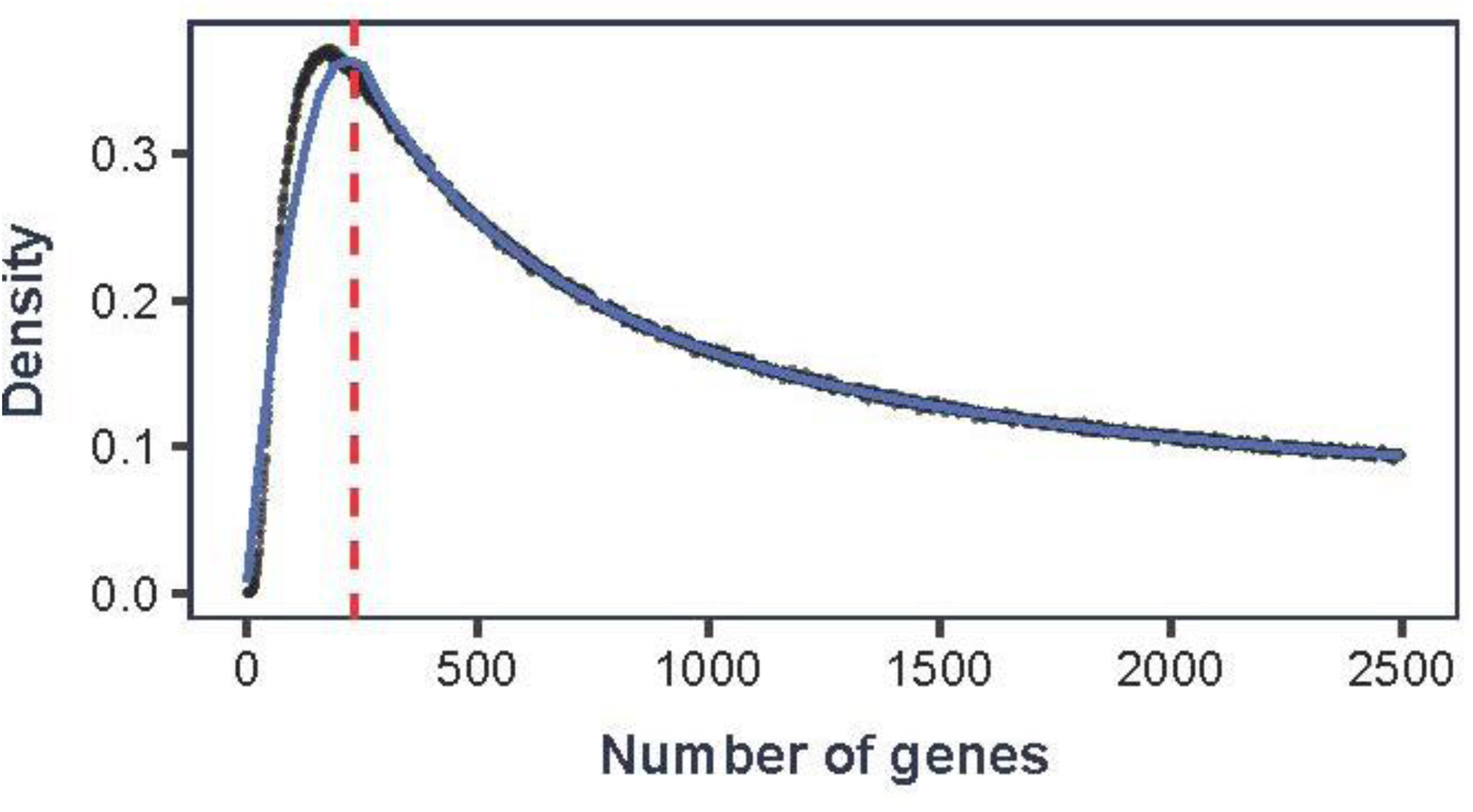
Maximum Likelihood Estimate (MLE) of number of CMS risk genes. For each number of possible risk genes between 1-2,500, we conducted 50,000 simulations to determine the number of risk genes that yielded the closest agreement between our observed and simulated data. This MLE method yields an estimate of 184 CMS risk genes (red vertical line). See Supplementary Methods.

**Figure S3.**
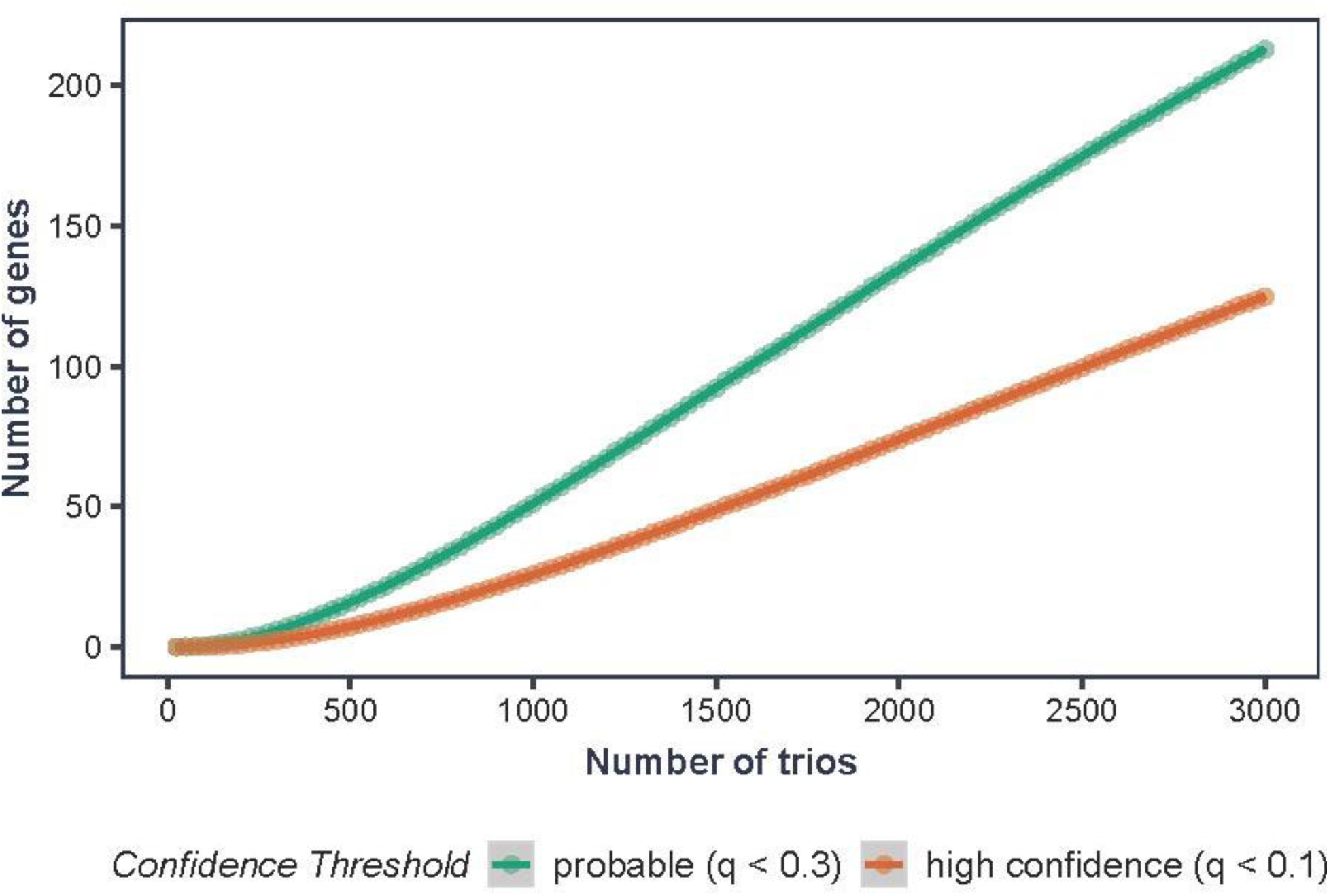
Gene discovery by number of trios sequenced. Using our estimate of 184 risk genes (based on the MLE method – see Main Text and Supplementary Methods), we estimated the number of probable (FDR q<0.3) and high-confidence (FDR q<0.1) risk genes that will be discovered as more CMS trios are sequenced. We performed 10,000 simulations at each cohort size from 25-3,000 trios, randomly generating variants and assigning to risk genes in agreement with the proportions seen in our data, then applying the TADA-Denovo algorithm. Based on these simulations, WES of 500 trios should find 16 probable and 7 high-confidence risk genes; 1000 trios should find 51 probable and 26 high-confidence risk genes.

**Figure S4.**
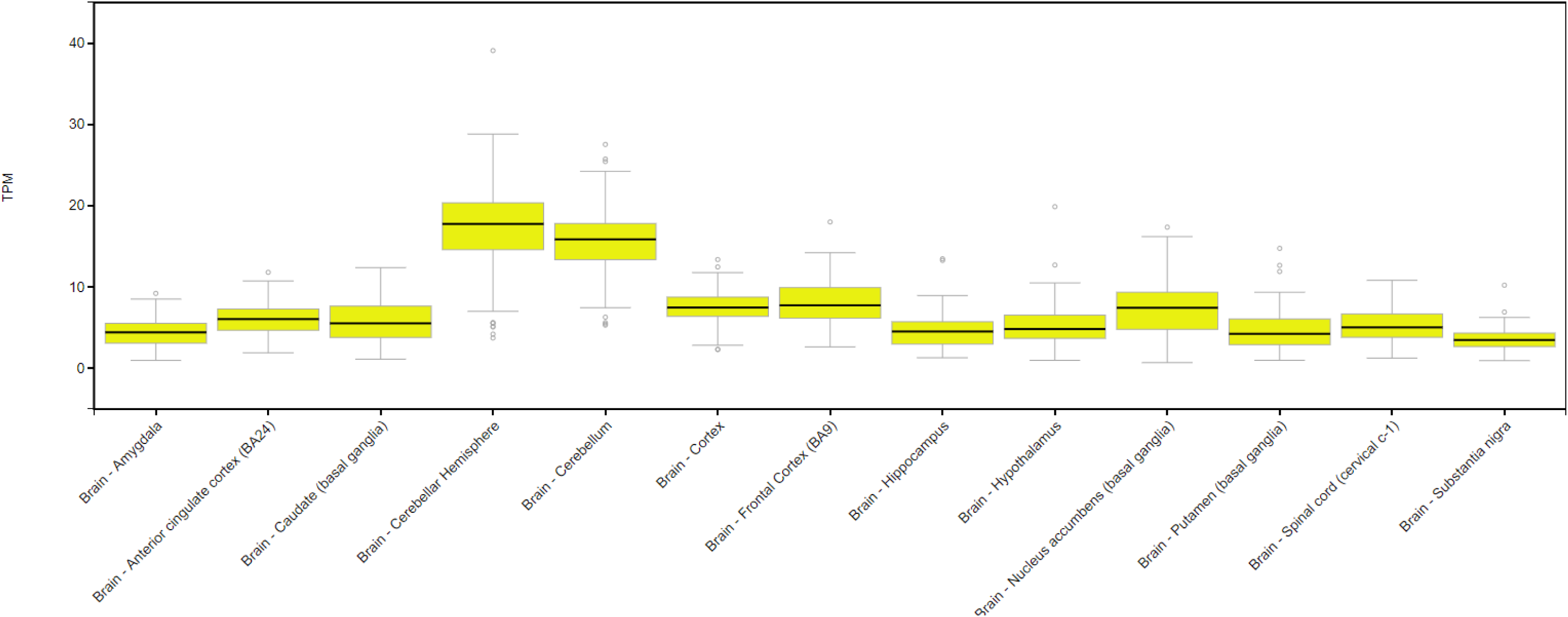
KDM5B brain expression levels. Brain expression by region for KDM5B. Data is from GTEx Analysis Release V7 (dbGaP Accession phs000424.v7.p2) (https://gtexportal.org/home/gene/KDM5B). Expression values are shown in Transcripts Per Million (TPM), calculated from a gene model with isoforms collapsed to a single gene. No other normalization steps have been applied. Box plots are shown as median, 25th, and 75^th^ percentiles. Points are displayed as outliers if they are above or below 1.5 times the interquartile range. Further details about expression quantification and samples can be found at https://gtexportal.org/home/documentationPage#AboutData.

**Figure S5.**
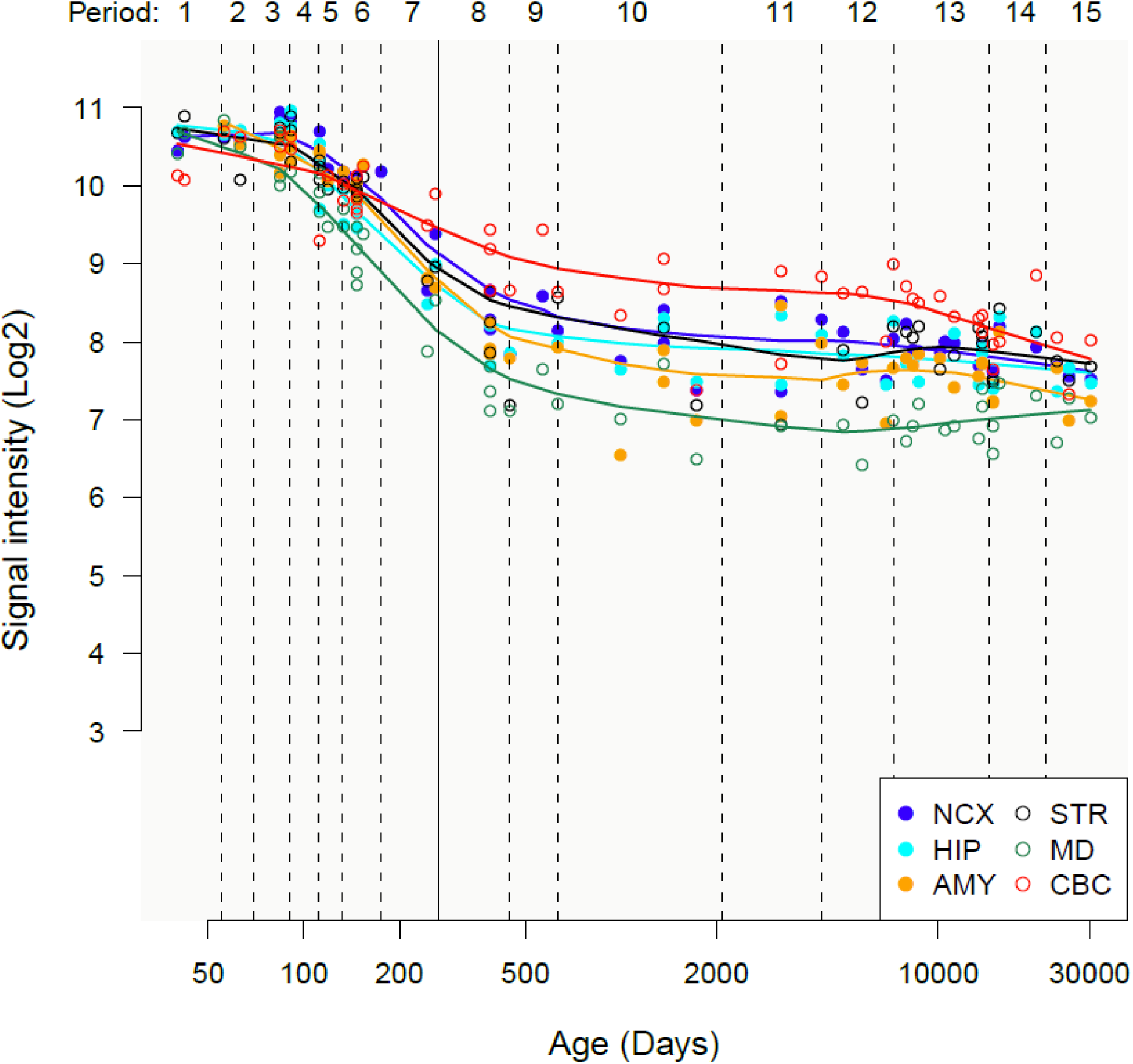
KDM5B spatiotemporal brain expression. Brain expression trajectories of *KDM5B* in the developing human brain. Expression data is from the Brainspan Consortium (brainspan.org, hbatlas.org), generated using the Affymetrix GeneChip Human Exon 1.0 ST Array platform. Vertical axis is the log2-transformed array signal intensity, which is proportional to transcript expression. A stringent threshold of ≥6 was required to meet criteria for brain expression. Horizontal axis represents periods of human development and adulthood as previously defined by Kang et al (2011). Birth begins period 8 and adolescence begins period 12. Brain regions are by color: neocortex (NCX), hippocampus (HIP), amygdala (AMY), striatum (STR), mediodorsal nucleus of the thalamus (MD), cerebellar cortex (CBC).

**Figure S6.**
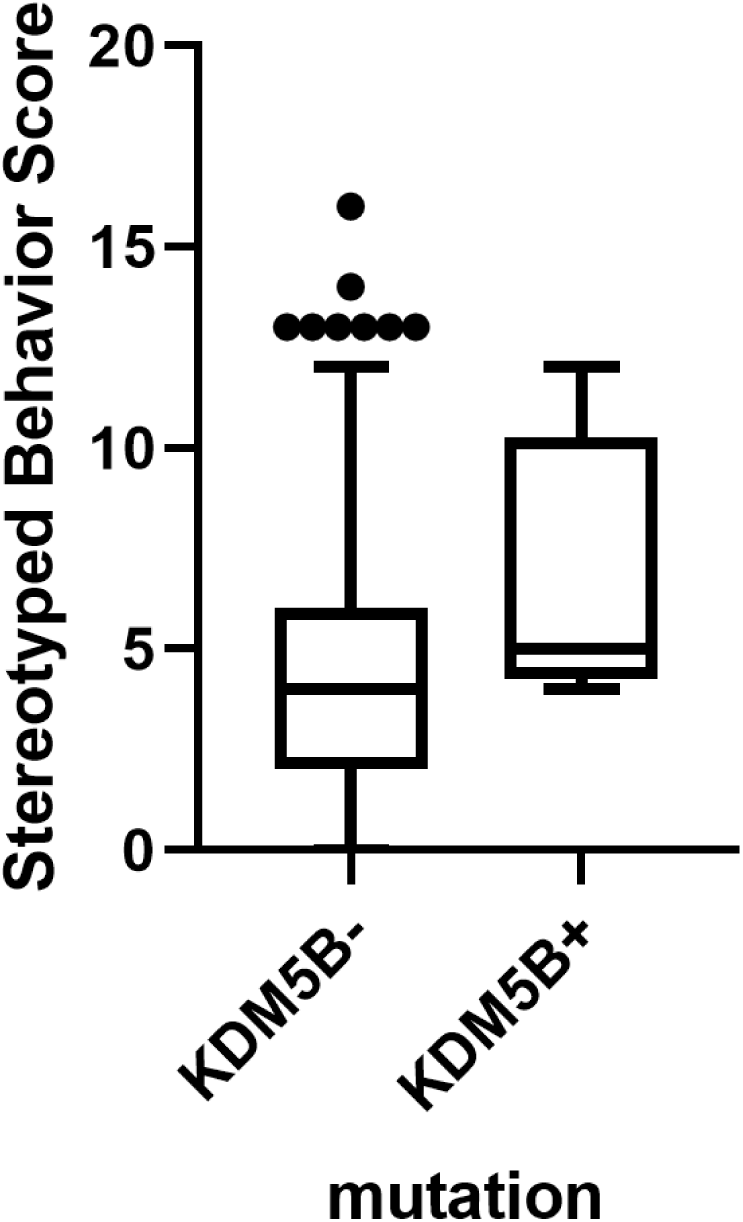
Stereotypy scores in Simons Simplex Collection ASD probands. Tukey box and whisker plot of Stereotyped Behavior Score (SBS) from the RBS-R (Repetitive Behavior Scale-Revised) in aged-matched Simons Simplex Collection ASD probands with (+, n=4) and without (-, n=364) de novo damaging mutations in *KDM5B*. Two-tailed Mann-Whitney test of ages (months) between groups: p=0.86. One-tailed Mann-Whitney test of SBS between groups: p=0.076.

## SUPPLEMENTARY TABLES

**Table S1 – Phenotype, exome sequencing metrics, and principal components analysis.**

*(see “TableS1.xlsx”)*

First tab contains individual-level sample information (columns A-I), including family ID, individual ID, phenotype, cohort, collection site, gender, capture platform, size of “callable exome”, and paternal age (years) at birth, where available. Column J lists reasons for any sample exclusions by quality control methods; “0” indicates that the sample was not excluded, and was included in subsequent analyses. Columns K-AF list individual sample sequencing metrics generated using PicardTools, and GATK DepthOfCoverage tools. Columns AG-AQ list individual sample sequencing metrics generated using PLINK/SEQ (i-stats; https://psychgen.u.hpc.mssm.edu/plinkseq/stats.shtml). Columns B, K-AQ were included in Principal Components Analysis (PCA). Third tab contains cohort-level metrics calculated using samples passing quality control. ±95% confidence intervals are given, when applicable. Fourth tab contains coordinates generated for each sample for the top 10 principal components following PCA. The code used to generate this data is included in Supplementary Methods. Using these coordinates, we removed trios with family members falling more than three standard deviations from the mean in any of the first five principal components; this information is contained in the fifth tab.

**Table S2 – Annotated de novo variants in CMS and controls.**

*(see “TableS2.xlsx”)*

Detailed information on all high confidence de novo variants in cases and controls. These variants were annotated using ANNOVAR, based on RefSeq hg19 gene definitions. Column descriptions are provided in a separate tab of this file. A third tab provides the number of each de novo variant type per sample.

**Table S3 – Gene-level de novo mutation rates, variant counts, and TADA-Denovo results.**

*(see “TableS3.xlsx”)*

First tab contains de novo mutation rates used to perform subsequent maximum likelihood estimation (MLE) and TADA-Denovo analyses. The following mutation rates are listed for each gene: overall (mut.rate), likely gene disrupting (lgd), predicted damaging missense (mis3), and all damaging (lgd + misD). These overall mutation rates were previously published (Ware et al., 2015) from unaffected parent-child trios. The code used to generate the mutation rate table is provided in Supplementary Methods. Second tab contains the input file for the TADA-Denovo algorithm. Gene-level expected mutation rates for LGD (“mut.cls1” column) and Mis-D variants (“mut.cls2” column) are listed, along with their respective observed mutation counts in our CMS data (“dn.cls1” and “dn.cls2”, respectively). Code for running TADA-Denovo is given in Supplementary Methods. Third tab contains the final output results from TADA-Denovo code provided in Supplementary Methods. One gene harboring more than one damaging de novo (LGD or Mis-D) variant in unrelated CMS families is highlighted in yellow (KDM5B). This gene exceeded the threshold for being considered a high confidence (qval < 0.1) risk gene.

**Table S4 – DNENRICH gene lists and results.**

*(see “TableS4.xlsx”)*

See Supplementary Methods for details of DNENRICH analysis and gene lists used. First tab contains input gene lists and information about their curation. Second tab contains the input mutation list for DNENRICH; each row represents a de novo damaging mutation in a CMS proband. Third tab contains results output from DNENRICH. Significantly enriched gene sets are highlighted.

**Table S5 – Exploratory pathway, gene ontology, and spatiotemporal analyses results.**

*(see “TableS5.xlsx”)*

Pathway and gene ontology results from ConsensusPathDB are in the first tab; p-values < 0.05 and corresponding q-values are shown. Specific Enrichment Analysis (SEA) exploring whether genes cluster within certain brain regions across development using Brainspan atlas data is in the second tab; p-values < 0.05 are highlighted in yellow. See Supplementary Methods for details of these analyses.

## REFERENCES

1. H. S. Singer, Stereotypic movement disorders. Handbook of clinical neurology 100, 631–639 (2011).

2. American Psychiatric Association, Diagnostic and statistical manual of mental disorders : DSM-5. (2013).

3. R. Militerni, C. Bravaccio, C. Falco, C. Fico, M. T. Palermo, Repetitive behaviors in autistic disorder. Eur Child Adolesc Psychiatry 11, 210–218 (2002).

4. S. L. Bishop, J. Richler, C. Lord, Association between restricted and repetitive behaviors and nonverbal IQ in children with autism spectrum disorders. Child Neuropsychol 12, 247–267 (2006).

5. S. Goldman et al., Motor stereotypies in children with autism and other developmental disorders. Dev Med Child Neurol 51, 30–38 (2009).

6. E. Honey, S. Leekam, M. Turner, H. McConachie, Repetitive behaviour and play in typically developing children and children with autism spectrum disorders. J Autism Dev Disord 37, 1107–1115 (2007).

7. M. Campbell et al., Stereotypies and tardive dyskinesia: abnormal movements in autistic children. Psychopharmacol Bull 26, 260–266 (1990).

8. J. W. Bodfish, F. J. Symons, D. E. Parker, M. H. Lewis, Varieties of repetitive behavior in autism: comparisons to mental retardation. J Autism Dev Disord 30, 237–243 (2000).

9. J. L. Matson, S. L. Kiely, J. W. Bamburg, The effect of stereotypies on adaptive skills as assessed with the DASH-II and Vineland Adaptive Behavior Scales. Res Dev Disabil 18, 471–476 (1997).

10. E. Gal, M. J. Dyck, A. Passmore, The relationship between stereotyped movements and self-injurious behavior in children with developmental or sensory disabilities. Res Dev Disabil 30, 342–352 (2009).

11. F. J. Symons, L. A. Sperry, P. L. Dropik, J. W. Bodfish, The early development of stereotypy and self-injury: a review of research methods. J Intellect Disabil Res 49, 144–158 (2005).

12. C. A. Doyle, C. J. McDougle, Pharmacologic treatments for the behavioral symptoms associated with autism spectrum disorders across the lifespan. Dialogues in clinical neuroscience 14, 263–279 (2012).

13. A. Tan, M. Salgado, S. Fahn, The characterization and outcome of stereotypical movements in nonautistic children. Mov Disord 12, 47–52 (1997).

14. E. M. Mahone, D. Bridges, C. Prahme, H. S. Singer, Repetitive arm and hand movements (complex motor stereotypies) in children. The Journal of pediatrics 145, 391–395 (2004).

15. S. Leekam et al., Repetitive behaviours in typically developing 2-year-olds. J Child Psychol Psychiatry 48, 1131–1138 (2007).

16. K. M. Harris, E. M. Mahone, H. S. Singer, Nonautistic motor stereotypies: clinical features and longitudinal follow-up. Pediatric neurology 38, 267–272 (2008).

17. L. G. Foster, Nervous habits and stereotyped behaviors in preschool children. J Am Acad Child Adolesc Psychiatry 37, 711–717 (1998).

18. N. Rafaeli-Mor, L. Foster, G. Berkson, Self-reported body-rocking and other habits in college students. Am J Ment Retard 104, 1–10 (1999).

19. R. MacDonald et al., Stereotypy in young children with autism and typically developing children. Res Dev Disabil 28, 266–277 (2007).

20. F. Sallustro, C. W. Atwell, Body rocking, head banging, and head rolling in normal children. The Journal of pediatrics 93, 704–708 (1978).

21. H. Tröster, Prevalence and functions of stereotyped behaviors in nonhandicapped children in residential care. J Abnorm Child Psychol 22, 79–97 (1994).

22. F. X. Castellanos, G. F. Ritchie, W. L. Marsh, J. L. Rapoport, DSM-IV stereotypic movement disorder: persistence of stereotypies of infancy in intellectually normal adolescents and adults. J Clin Psychiatry 57, 116–122 (1996).

23. G. T. Baranek, Autism during infancy: a retrospective video analysis of sensory-motor and social behaviors at 9-12 months of age. J Autism Dev Disord 29, 213–224 (1999).

24. E. Werner, G. Dawson, Validation of the phenomenon of autistic regression using home videotapes. Arch Gen Psychiatry 62, 889–895 (2005).

25. A. Loh et al., Stereotyped motor behaviors associated with autism in high-risk infants: a pilot videotape analysis of a sibling sample. J Autism Dev Disord 37, 25–36 (2007).

26. H. S. Singer, Motor stereotypies. Seminars in pediatric neurology 16, 77–81 (2009).

27. L. G. Foster, Nervous habits and stereotyped behaviors in preschool children. J Am Acad Child Adolesc Psychiatry 37, 711–717 (1998).

28. C. Oakley, E. M. Mahone, C. Morris-Berry, T. Kline, H. S. Singer, Primary complex motor stereotypies in older children and adolescents: clinical features and longitudinal follow-up. Pediatric neurology 52, 398–403.e391 (2015).

29. J. M. Miller, H. S. Singer, D. D. Bridges, H. R. Waranch, Behavioral therapy for treatment of stereotypic movements in nonautistic children. Journal of child neurology 21, 119–125 (2006).

30. M. W. Specht et al., Efficacy of parent-delivered behavioral therapy for primary complex motor stereotypies. Dev Med Child Neurol 59, 168–173 (2017).

31. S. Gao, H. S. Singer, Complex motor stereotypies: an evolving neurobiological concept. Future Neurology 8, 273–285 (2013).

32. J. C. Carter, G. T. Capone, W. E. Kaufmann, Neuroanatomic correlates of autism and stereotypy in children with Down syndrome. Neuroreport 19, 653–656 (2008).

33. A. E. Kelley, C. G. Lang, A. M. Gauthier, Induction of oral stereotypy following amphetamine microinjection into a discrete subregion of the striatum. Psychopharmacology (Berl) 95, 556–559 (1988).

34. W. R. Kates, D. C. Lanham, H. S. Singer, Frontal white matter reductions in healthy males with complex stereotypies. Pediatric neurology 32, 109–112 (2005).

35. S. Sato, T. Hashimoto, A. Nakamura, S. Ikeda, Stereotyped stepping associated with lesions in the bilateral medial frontoparietal cortices. Neurology 57, 711–713 (2001).

36. D. M. Maraganore, A. J. Lees, C. D. Marsden, Complex stereotypies after right putaminal infarction: a case report. Mov Disord 6, 358–361 (1991).

37. H. S. Singer, Motor control, habits, complex motor stereotypies, and Tourette syndrome. Annals of the New York Academy of Sciences 1304, 22–31 (2013).

38. E. M. Mahone et al., Anomalous Putamen Volume in Children With Complex Motor Stereotypies. Pediatric neurology 65, 59–63 (2016).

39. A. H. Evans et al., Punding in Parkinson’s disease: its relation to the dopamine dysregulation syndrome. Mov Disord 19, 397–405 (2004).

40. M. H. Lewis et al., Plasma HVA in adults with mental retardation and stereotyped behavior: biochemical evidence for a dopamine deficiency model. Am J Ment Retard 100, 413–418 (1996).

41. A. D. Harris et al., GABA and Glutamate in Children with Primary Complex Motor Stereotypies: An 1H-MRS Study at 7T. AJNR. American journal of neuroradiology 37, 552–557 (2016).

42. K. Dworzynski, F. Happé, P. Bolton, A. Ronald, Relationship between symptom domains in autism spectrum disorders: a population based twin study. J Autism Dev Disord 39, 1197–1210 (2009).

43. R. D. Freeman, A. Soltanifar, S. Baer, Stereotypic movement disorder: easily missed. Dev Med Child Neurol 52, 733–738 (2010).

44. F. K. Satterstrom et al., Large-scale exome sequencing study implicates both developmental and functional changes in the neurobiology of autism. bioRxiv 10.1101/484113, 484113 (2019).

45. C. Cappi et al., De novo damaging coding mutations are strongly associated with obsessive-compulsive disorder and overlap with autism. bioRxiv 10.1101/127712, 127712 (2017).

46. S. Wang et al., De Novo Sequence and Copy Number Variants Are Strongly Associated with Tourette Disorder and Implicate Cell Polarity in Pathogenesis. Cell reports 24, 3441–3454.e3412 (2018).

47. G. D. Fischbach, C. Lord, The Simons Simplex Collection: a resource for identification of autism genetic risk factors. Neuron 68, 192–195 (2010).

48. A. McKenna et al., The Genome Analysis Toolkit: a MapReduce framework for analyzing next-generation DNA sequencing data. Genome Res 20, 1297–1303 (2010).

49. C. Cappi et al., De novo damaging coding mutations are strongly associated with obsessive-compulsive disorder and overlap with autism. bioRxiv 10.1101/127712 (2017).

50. K. Wang, M. Li, H. Hakonarson, ANNOVAR: functional annotation of genetic variants from high-throughput sequencing data. Nucleic Acids Res 38, e164 (2010).

51. M. Lek et al., Analysis of protein-coding genetic variation in 60,706 humans. Nature 536, 285–291 (2016).

52. A. J. Willsey et al., De Novo Coding Variants Are Strongly Associated with Tourette Disorder. Neuron 94, 486–499 e489 (2017).

53. X. He et al., Integrated model of de novo and inherited genetic variants yields greater power to identify risk genes. PLoS Genet 9, e1003671 (2013).

54. S. J. Sanders et al., Insights into Autism Spectrum Disorder Genomic Architecture and Biology from 71 Risk Loci. Neuron 87, 1215–1233 (2015).

55. J. Homsy et al., De novo mutations in congenital heart disease with neurodevelopmental and other congenital anomalies. Science 350, 1262–1266 (2015).

56. M. Fromer et al., De novo mutations in schizophrenia implicate synaptic networks. Nature 506, 179–184 (2014).

57. R. Herwig, C. Hardt, M. Lienhard, A. Kamburov, Analyzing and interpreting genome data at the network level with ConsensusPathDB. Nat Protoc 11, 1889–1907 (2016).

58. H. J. Kang et al., Spatio-temporal transcriptome of the human brain. Nature 478, 483–489 (2011).

59. J. D. Dougherty, E. F. Schmidt, M. Nakajima, N. Heintz, Analytical approaches to RNA profiling data for the identification of genes enriched in specific cells. Nucleic Acids Res 38, 4218–4230 (2010).

60. R. Cantor, Autism endophenotypes and quantitative trait loci. Autism Spectrum Disorders. Oxford University Press: New York, 690–704 (2011).

61. R. M. Cantor et al., ASD restricted and repetitive behaviors associated at 17q21.33: genes prioritized by expression in fetal brains. Mol Psychiatry 23, 993–1000 (2018).

62. P. B. Rasmussen, P. Staller, The KDM5 family of histone demethylases as targets in oncology drug discovery. Epigenomics 6, 277–286 (2014).

63. X. Li et al., Histone demethylase KDM5B is a key regulator of genome stability. Proceedings of the National Academy of Sciences 111, 7096–7101 (2014).

64. B. L. Kidder, G. Hu, K. Zhao, KDM5B focuses H3K4 methylation near promoters and enhancers during embryonic stem cell self-renewal and differentiation. Genome Biology 15, R32 (2014).

65. S. Zaidi et al., De novo mutations in histone modifying genes in congenital heart disease. Nature 498, 220–223 (2013).

66. J. R. Horton et al., Characterization of a Linked Jumonji Domain of the KDM5/JARID1 Family of Histone H3 Lysine 4 Demethylases. The Journal of biological chemistry 291, 2631–2646 (2016).

## References

1. J. M. Miller, H. S. Singer, D. D. Bridges, H. R. Waranch, Behavioral therapy for treatment of stereotypic movements in nonautistic children. Journal of child neurology 21, 119–125 (2006).

2. S. Ehlers, C. Gillberg, L. Wing, A screening questionnaire for Asperger syndrome and other high-functioning autism spectrum disorders in school age children. J Autism Dev Disord 29, 129–141 (1999).

3. J. S. March, J. D. Parker, K. Sullivan, P. Stallings, C. K. Conners, The Multidimensional Anxiety Scale for Children (MASC): factor structure, reliability, and validity. J Am Acad Child Adolesc Psychiatry 36, 554–565 (1997).

4. G. J. DuPaul, T. J. Power, A. D. Anastopoulos, R. Reid, ADHD Rating Scale—IV: Checklists, norms, and clinical interpretation (Guilford Press, 1998).

5. L. Scahill et al., Children’s Yale-Brown Obsessive Compulsive Scale: reliability and validity. J Am Acad Child Adolesc Psychiatry 36, 844–852 (1997).

6. J. W. Bodfish, F. J. Symons, D. E. Parker, M. H. Lewis, Varieties of repetitive behavior in autism: comparisons to mental retardation. J Autism Dev Disord 30, 237–243 (2000).

7. J. N. Constantino (2005) Social Responsiveness Scale (SRS). (Western Psychological Services, Los Angeles, CA).

8. A. McKenna et al., The Genome Analysis Toolkit: a MapReduce framework for analyzing next-generation DNA sequencing data. Genome Res 20, 1297–1303 (2010).

9. C. Cappi et al., De novo damaging coding mutations are strongly associated with obsessive-compulsive disorder and overlap with autism. bioRxiv 10.1101/127712 (2017).

10. H. Li, R. Durbin, Fast and accurate long-read alignment with Burrows-Wheeler transform. Bioinformatics 26, 589–595 (2010).

11. K. Wang, M. Li, H. Hakonarson, ANNOVAR: functional annotation of genetic variants from high-throughput sequencing data. Nucleic Acids Res 38, e164 (2010).

12. A. Manichaikul et al., Robust relationship inference in genome-wide association studies. *Bioinformatics (Oxford*, England) 26, 2867–2873 (2010).

13. P. Danecek et al., The variant call format and VCFtools. Bioinformatics (Oxford, England) 27, 2156–2158 (2011).

14. M. Fromer et al., De novo mutations in schizophrenia implicate synaptic networks. Nature 506, 179–184 (2014).

15. C. Cappi et al., De novo damaging coding mutations are strongly associated with obsessive-compulsive disorder and overlap with autism. bioRxiv 10.1101/127712, 127712 (2017).

16. C. Cappi et al., Whole-exome sequencing in obsessive-compulsive disorder identifies rare mutations in immunological and neurodevelopmental pathways. Transl Psychiatry 6, e764 (2016).

17. M. Lek et al., Analysis of protein-coding genetic variation in 60,706 humans. Nature 536, 285–291 (2016).

18. J. S. Ware, K. E. Samocha, J. Homsy, M. J. Daly, Interpreting de novo Variation in Human Disease Using denovolyzeR. Current protocols in human genetics / editorial board, Jonathan L. Haines … [et al.] 87, 7.25.21–15 (2015).

19. X. He et al., Integrated model of de novo and inherited genetic variants yields greater power to identify risk genes. PLoS Genet 9, e1003671 (2013).

20. S. J. Sanders et al., Insights into Autism Spectrum Disorder Genomic Architecture and Biology from 71 Risk Loci. Neuron 87, 1215–1233 (2015).

21. A. J. Willsey et al., De Novo Coding Variants Are Strongly Associated with Tourette Disorder. Neuron 94, 486–499 e489 (2017).

22. J. Homsy et al., De novo mutations in congenital heart disease with neurodevelopmental and other congenital anomalies. Science 350, 1262–1266 (2015).

23. S. Shohat, E. Ben-David, S. Shifman, Varying Intolerance of Gene Pathways to Mutational Classes Explain Genetic Convergence across Neuropsychiatric Disorders. Cell reports 18, 2217–2227 (2017).

24. S. L. Girard et al., Increased exonic de novo mutation rate in individuals with schizophrenia. Nat Genet 43, 860–863 (2011).

25. S. Gulsuner et al., Spatial and temporal mapping of de novo mutations in schizophrenia to a fetal prefrontal cortical network. Cell 154, 518–529 (2013).

26. S. E. McCarthy et al., De novo mutations in schizophrenia implicate chromatin remodeling and support a genetic overlap with autism and intellectual disability. Mol Psychiatry 19, 652–658 (2014).

27. B. Xu et al., Exome sequencing supports a de novo mutational paradigm for schizophrenia. Nat Genet 43, 864–868 (2011).

28. J. de Ligt et al., Diagnostic exome sequencing in persons with severe intellectual disability. The New England journal of medicine 367, 1921–1929 (2012).

29. C. Gilissen et al., Genome sequencing identifies major causes of severe intellectual disability. Nature 511, 344–347 (2014).

30. F. F. Hamdan et al., De novo mutations in moderate or severe intellectual disability. PLoS Genet 10, e1004772 (2014).

31. A. Rauch et al., Range of genetic mutations associated with severe non-syndromic sporadic intellectual disability: an exome sequencing study. Lancet 380, 1674–1682 (2012).

32. Deciphering Developmental Disorders Study, Prevalence and architecture of de novo mutations in developmental disorders. Nature 10.1038/nature21062 (2017).

33. S. Wang et al., De Novo Sequence and Copy Number Variants Are Strongly Associated with Tourette Disorder and Implicate Cell Polarity in Pathogenesis. Cell reports 24, 3441–3454.e3412 (2018).

34. F. K. Satterstrom et al., ASD and ADHD have a similar burden of rare protein-truncating variants. bioRxiv 10.1101/277707, 277707 (2018).

35. R. Herwig, C. Hardt, M. Lienhard, A. Kamburov, Analyzing and interpreting genome data at the network level with ConsensusPathDB. Nat Protoc 11, 1889–1907 (2016).

36. H. J. Kang et al., Spatio-temporal transcriptome of the human brain. Nature 478, 483–489 (2011).

37. J. D. Dougherty, E. F. Schmidt, M. Nakajima, N. Heintz, Analytical approaches to RNA profiling data for the identification of genes enriched in specific cells. Nucleic Acids Res 38, 4218–4230 (2010).

